# Crystal structures of glycoprotein D of equine alphaherpesviruses reveal potential binding sites to the entry receptor MHC-I

**DOI:** 10.1101/2022.06.10.495596

**Authors:** Viviane Kremling, Bernhard Loll, Szymon Pach, Ismail Dahmani, Christoph Weise, Gerhard Wolber, Salvatore Chiantia, Markus C. Wahl, Nikolaus Osterrieder, Walid Azab

**Affiliations:** Institut für Virologie, Robert von Ostertag-Haus, Zentrum für Infektionsmedizin, Freie Universität Berlin, Berlin, Germany; Laboratory of Structural Biochemistry, Freie Universität Berlin, Berlin, Germany; Institute of Pharmacy (Pharmaceutical Chemistry), Freie Universität Berlin, Berlin, Germany; Universität Potsdam, Institut für Biochemie und Biologie, Potsdam, Brandenburg, Germany; BioSupraMol Core Facility, Bio-Mass Spectrometry, Freie Universität Berlin, Berlin, Germany; Helmholtz-Zentrum Berlin für Materialien und Energie, Macromolecular Crystallography, Berlin, Germany; Department of Infectious Diseases and Public Health, City University of Hong Kong, Kowloon Tong, Hong Kong

## Abstract

Cell entry of most alphaherpesviruses is mediated by the binding of glycoprotein D (gD) to different cell surface receptors. Equine herpesvirus type 1 (EHV-1) and EHV-4 gDs interact with equine major histocompatibility complex I (MHC-I) to initiate entry into equine cells. We have characterized the gD-MHC-I interaction by solving the crystal structures of EHV-1 and EHV-4 gDs (gD1, gD4), performing protein-protein docking simulations, surface plasmon resonance (SPR) analysis, and biological assays. The structures of gD1 and gD4 revealed the existence of a common V-set immunoglobulin-like (IgV-like) core comparable to those of other gD homologs. Molecular modeling yielded plausible binding hypotheses and identified key residues (F213 and D261) that are important for virus binding. Altering the key residues resulted in impaired virus growth in cells, which highlights the important role of these residues in the gD-MHC-I interaction. Taken together, our results add to our understanding of the initial herpesvirus-cell interactions and will contribute to the targeted design of antiviral drugs and vaccine development.

**Author summary:** Equine herpesvirus type 1 (EHV-1) and type 4 (EHV-4) are endemic in horses and cause great suffering as well as substantial economic losses to the equine industry. Current vaccines do not prevent infections and treatment is difficult. A prerequisite for vaccine and drug development is an in-depth understanding of the virus replication cycle, especially the virus entry process in order to block the infection at early stages. Entry of alphaherpesviruses into the host cell is mediated by a set of virus envelope glycoproteins including glycoprotein D (gD) that triggers the internalization of the virus particle. The structure of gD and the interaction with the entry receptor equine major histocompatibility complex class I (MHC-I) remains elusive. Here, we solved the crystal structures of gD1 and gD4 that allowed us to model virus-receptor interaction and to determine the key residues for virus entry. Alterations of these key residues impaired virus growth in cell culture. The overall structure of gD1 and gD4 shows classical features of other alphaherpesvirus gDs making it possible to gain further insights into human pathogens as well.

## Introduction

One of the most essential steps for virus replication is the entry process into host cells. In herpesviruses, more specifically alphaherpesviruses, cell entry is a complex multistep process that requires a stepwise contribution of five out of twelve envelope glycoproteins, namely glycoprotein B (gB), gC, gD, and the heterodimer gH/gL [1]. Of these, gD is the (main) receptor-binding protein that interacts with the cell receptors and triggers the subsequent fusion process with cell membrane and/or uptake by endocytosis [2–4].

Equine herpesvirus type 1 (EHV-1) and EHV-4 use equine major histocompatibility complex class I (MHC-I) as an entry receptor, however, no details of the molecular binding mode are available [5–7]. Only few other viruses are known to utilize MHC molecules as binding receptors but not as entry receptors. Coxsackievirus A9 requires MHC-I and GRP78 as co-receptors for virus internalization [8], Simian virus 40 (SV40) binds to cellular MHC-I, however, MHC-I does not mediate virus entry [9, 10]. The fiber knob of Adenovirus type 5 (AdV-5) binds to the α2 region of human leukocyte antigen (HLA) [11],and the functional gD homolog gp42 in Epstein-Barr Virus (EBV) binds to MHC-II to activate membrane fusion [12].

MHC-I seems to be an unlikely receptor for viral entry since it is present on all somatic cells [13] and therefore does not confer tissue specificity. Additionally, it is one of the most polymorphic mammalian proteins with 10 to 25% difference in the amino acid sequence [14, 15]. Typically, MHC-I plays a crucial role in the adaptive immunity by presenting proteolytically processed intracellular proteins on the cell surface to T-cells and natural killer cells [16]. In case of an infected cell, virus-derived peptides are presented and the recognition by T-cell receptor (TCR) initiates an immune response [17]. Although utilized by EHV-1 and EHV-4 as entry receptors, not all MHC-I genes support the entry of both viruses [5,7,18]. Interestingly, the residue A173 in the α2 region of MHC-I seems to be necessary but not enough to trigger virus entry [7,18,19].

EHV-1 and EHV-4 are important pathogens that cause great suffering in *Equidae* and other mammals and result in huge economic losses to the equine industry [20]. Efforts have been made to find efficient vaccines against both viruses [21]. However, the protection is usually limited in time and efficacy; frequent outbreaks occur also in vaccinated horses [22–25]. Here, we present crystal structures of free gD1 and gD4, which show a similar fold as other gD proteins from related viruses such as herpes simplex virus type 1 (HSV-1) (PDB-ID 2C36, [26], pseudorabies virus (PrV, PDB-ID 5X5V, 27), and bovine herpesvirus type 1 (BoHV-1, PDB-ID 6LS9, [27]. We further measured dissociation constants (in a micromolar range) for recombinant gD1/gD4 and C-terminally truncated gD4 binding to equine MHC-I by surface plasmon resonance spectroscopy (SPR). No increased binding affinity was observed for the truncated protein as was the case for gD of HSV-1, HSV-2, and PrV [28, 29], suggesting a structurally different mode of binding during entry into host cells. Cell culture assays showed that recombinant gD1 and gD4 as well as truncated gD4 can inhibit viral replication *in vitro*, where again the truncated version did not perform better than the full-length protein. The crystal structures were further used for *in silico* docking analyses to equine MHC-I. Based on these docking positions, viral mutants with point mutations at position F213 or D261 were produced and displayed significant growth impairments which support the proposed mode of binding of gD1 and gD4 to MHC-I.

## Results

### Crystal structure of unbound EHV-1 and EHV-4 gD

Recombinant gD1 and gD4 lacking the transmembrane region were produced in insect cells, purified by affinity and size exclusion chromatography and used for crystallization experiments (Figure S 1 and Figure S 2). To evaluate the correct identity, sequence, and molecular mass of gD1, gD4, and equine MHC-I, mass spectrometry (MS) analysis was conducted. Recombinant equine MHC-I 3.1 (Eqca-1*00101) including a peptide (SDYVKVSNI) linked to the β2-microglobulin (β2m) region was produced in insect cells as well and purified in the same manner as gD1 and gD4.

To this point, molecular masses of gD1 and gD4 were determined only by SDS-PAGE and Western blotting. However, these techniques are known to often lead to an overestimation of the molecular mass [30]. Here we employed matrix-assisted laser desorption ionization-time of flight mass spectrometry (MALDI-TOF-MS) to analyze diluted recombinant protein. The proteins gD1, gD4 and the MHC-I α-chain contain a Tobacco Etch Virus (TEV) cleavage site and a His_6_-tag (ENLYFQG-H_6_) which contribute approximately 1675 Da to the molecular weight of the molecules (calculated with https://web.expasy.org/peptide_mass). Additionally, the residues EF (approximately 300 Da) originating from the *Eco-RI* restriction site, were detected by in-source-decay (ISD) in the recombinant proteins. A size of approximately 43078 Da for gD1, 43761 Da for gD4, 37779 Da for the α-chain of MHC-I and 13241 Da for β2m with its linker and attached peptide was determined (Figure 1 A-C). Excluding the molecular weight of the TEV cleavage site and the His_6_-tag (1675 Da), this translates into a molecular weight of 41403 and 42086 Da for soluble gD1 and gD4, respectively (Table 1) and implies an approximate molecular weight of 49345 Da for the recombinant MHC-I molecule consisting of α-chain (36100 Da) and β2m with linker and peptide (13240 Da), and 48500 Da without the linker.

**Figure 1:**
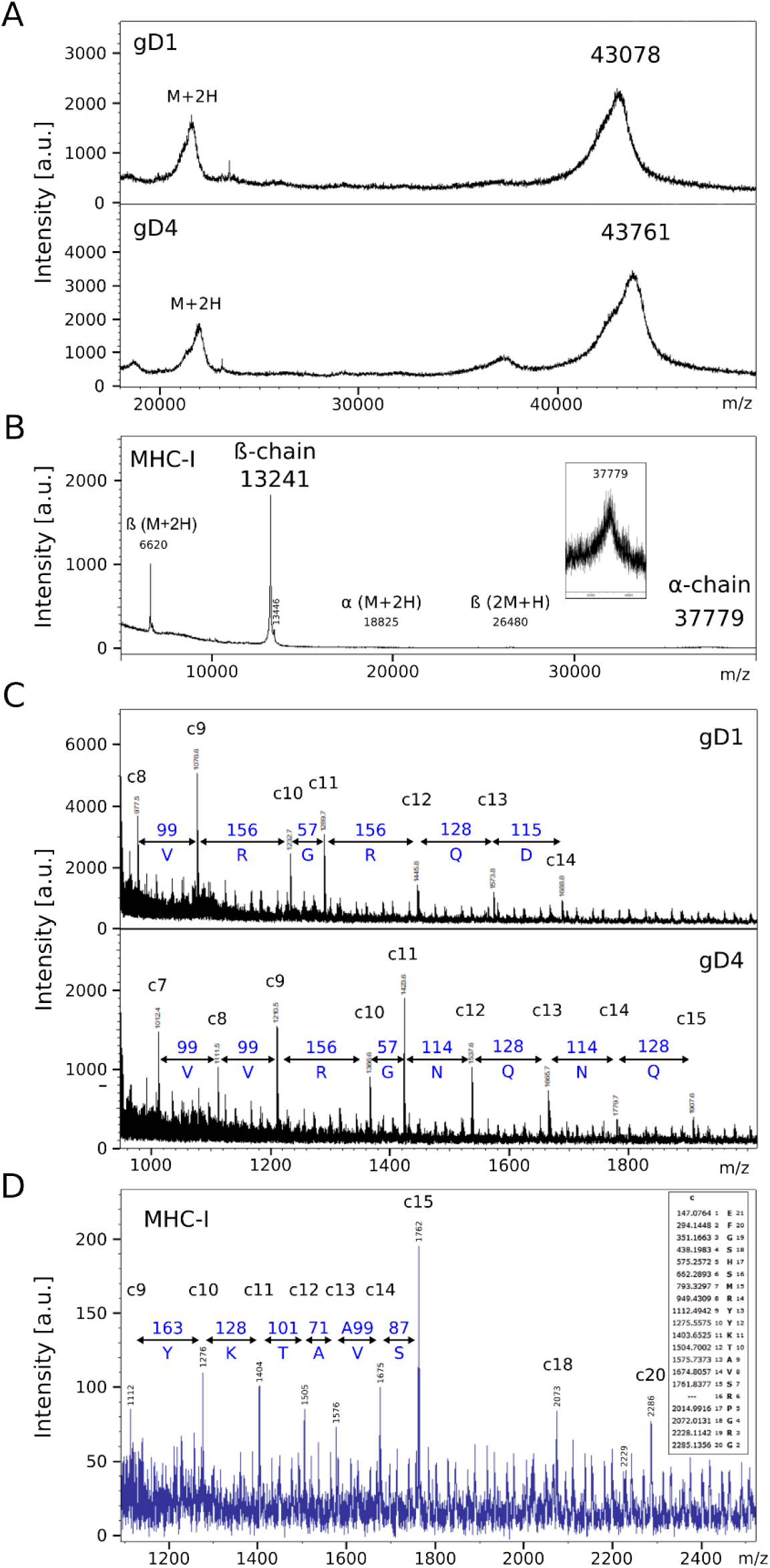
Mass spectrometric analysis of gD1 and gD4. (A) Intact protein mass analysis of recombinant gD1 (top) and gD4 (bottom) including N-terminal residues (EF: glutamic acid and phenylalanine) from *Eco*RI restriction site, TEV cleavage site and His_6_-tag on sinapinic acid (SA) matrix. (B) Intact protein mass analysis of recombinant MHC-I complex comprised of β2m and MHC-I-α-chain (insert zoom) including the same additional residues as gD1 and gD4. (C) In-source decay (ISD) spectra of recombinant gD1 (top), gD4 (bottom), and (D) MHC-I to ascertain the correct N-termini, insert: Theoretical c-ion series including the N-terminal EF extension.

**Table 1:**
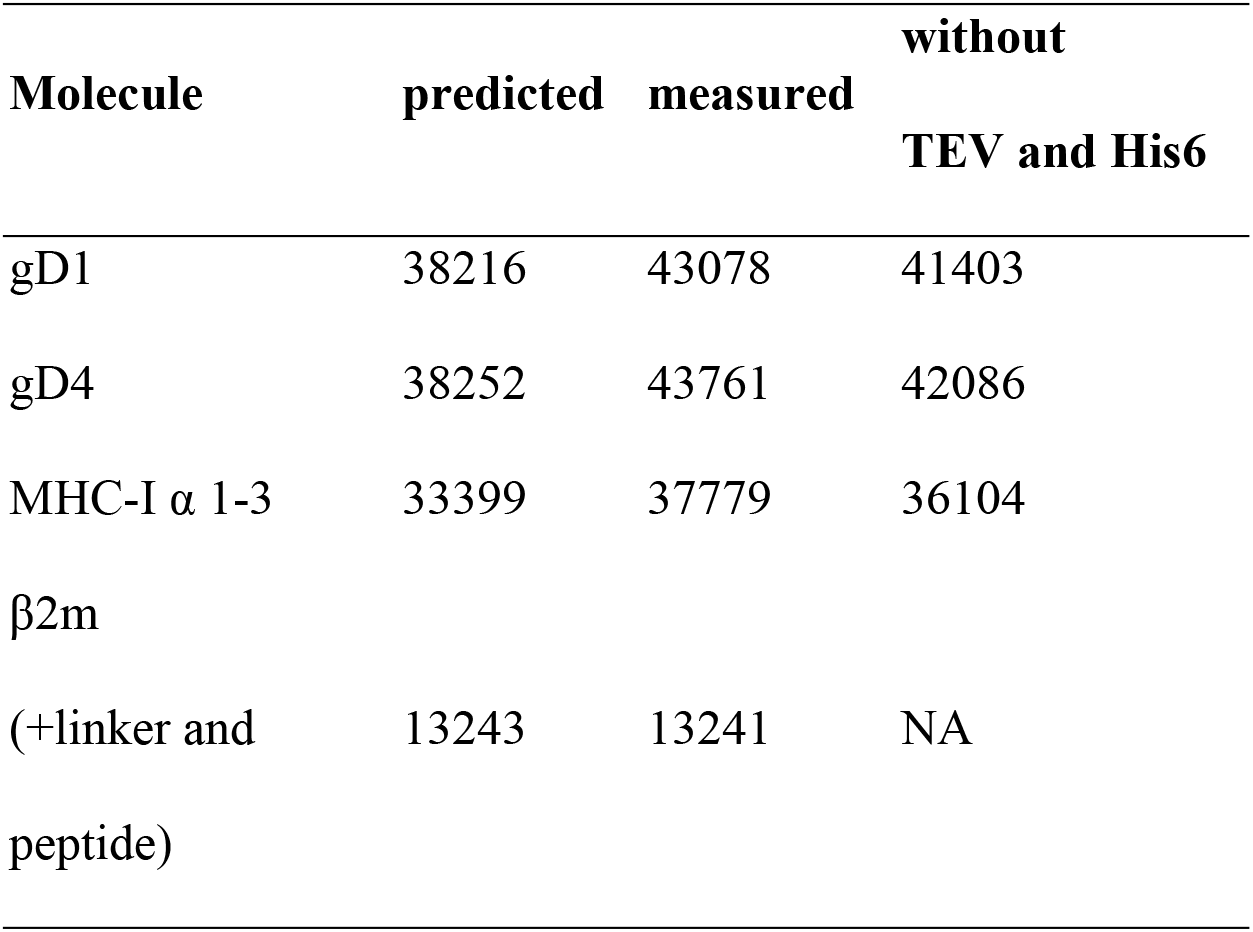
Predicted and measured molecular mass in Da of recombinant gD1, gD4 (aa 31-349), and equine MHC-I 3.1 α and β2m region with the uncleaved TEV site and His_6_-tag. Theoretical masses were calculated using https://web.expasy.org/peptide_mass/ and the actual mass determined by matrix-assisted laser desorption ionization-time of flight mass spectrometry (MALDI-TOF-MS). Post-translational modifications like glycosylations account for the discrepancies between the predictions and measurements. NA: not applicable.

The difference between predicted and observed molecular masses is due to post translational modifications (PTMs) such as glycosylations which contribute approximately 4000-5000 Da (Table 1). Further analysis of recombinant gD1, gD4, and MHC-I by ISD and tandem mass spectrometry (MS/MS) of in-gel digested Coomassie-stained proteins confirmed protein identities and presence of the correct N- and C-termini of gD1 and gD4 and N-terminus of MHC-I α-chain (Figure S 4 A, B, C).

A 2.45 Å resolution diffraction data set was collected for a gD1 crystal, and the gD1 structure (Figure 2 A) was determined using the structure coordinates of HSV-1 gD (protein data bank PDB-ID 2C36) for molecular replacement and refined to an R_work_ of 20.3% and R_free_ of 25.7% (Table 2, PDB-ID 6SQJ). The crystal structure of gD1 contains two gD molecules per asymmetric unit. In solution only the monomeric form was observed by size exclusion chromatography (SEC)-multi-angle static light scattering (MALS) (Figure S 5). Two ions interpreted as magnesium are trapped between the molecules and coordinated by residues E242 and D261 of both protein chains together with water molecules. The residues E32 to R38 and N281 to T348 could not be modeled due to a lack of electron density. N-linked glycosylations are visible at the predicted sites N20 and N28 [31] which are conserved between gD1 and gD4 but not in gDs of other alphaherpesviruses.

**Figure 2:**
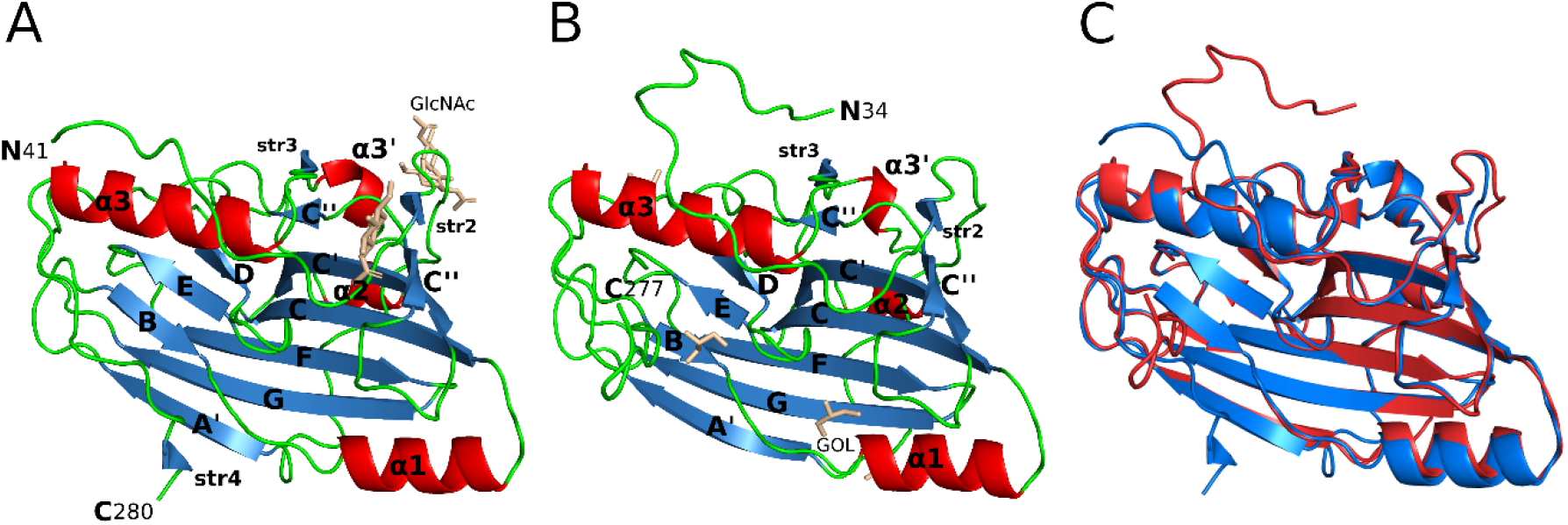
Crystal structure comparison of gD1 and gD4. Cartoon representation of (A) gD1 (2.45 Å resolution, PDB-ID: 6SQJ) and (B) gD4 (1.9 Å resolution, PDB-ID: 6TM8) monomer crystal structures. Molecule orientation is identical and secondary structures were assigned with dssp [72]. Helices are displayed in red, sheets in blue, and loops in green. N-acetylglucosamine (GlcNAc) and glycerol (GOL) molecules are shown in stick representation in beige. (C) Superposition of the crystal structures of gD1 (blue, PDB-ID 6SQJ) and gD4 (red, PDB-ID 6TM8). GlcNAc and glycerol molecules are not shown.

**Table 2:** Crystallographic data collection and model refinement statistics.

|  | **gD1** | **gD4** |
| --- | --- | --- |
| PDB-ID | **6SQJ** | **6TM8** |
| **Data collection** |  |  |
| Wavelength [Å] | 1.0332 | 0.91841 |
| Temperature [K] | 100 | 100 |
| Space group | P2<sub>1</sub>2<sub>1</sub>2<sub>1</sub> | P2<sub>1</sub>2<sub>1</sub>2<sub>1</sub> |
| **Unit cell parameters** |  |  |
| a, b, c [Å] | 71.9; 94.5; 101.3 | 73.1; 59.6; 69.7 |
| $\alpha=\beta=\gamma$ [°] | 90 | 90 |
| Resolution range [Å] | 50.00 - 2.24 (2.38 - 2.24)<sup>a</sup> | 50.00 - 1.90 (2.01 - 1.90) |
| Reflections<sup>a</sup> | 218509 (33751)<sup>a</sup> | 138685 (10835) |
| Unique reflections<sup>a</sup> | 33402 (5140) | 23671 (1810) |
| Completeness [%]<sup>a</sup> | 99.1 (95.8) | 95.6 (78.2) |
| Multiplicity<sup>a</sup> | 6.5 (6.6) | 5.9 (3.5) |
| **Data quality<sup>a</sup>** |  |  |
| I/$\sigma$(I)<sup>a</sup> | 11.71 (0.92) | 8.96 (0.96) |
| R<sub>meas</sub> [%]<sup>a</sup> | 13.5 (199) | 17.4 (126.8) |
| CC<sub>1/2</sub><sup>a</sup> | 99.8 (58.6) | 99.5 (40.9) |
| Wilson B factor [Å<sup>2</sup>] | 53.3 | 32 |
| **Refinement gD1 gD4** |  |  |
| Resolution range [Å]<sup>a</sup> | 50.00 - 2.24 (2.33 - 2.24) | 50.00 - 1.90 (2.01 - 1.90) |
| Reflections<sup>a</sup> | 33399 (3181) | 23642 (1792) |
| Test set (5%)<sup>a</sup> | 1669 (159) | 1182 (89) |
| R<sub>work</sub> [%]<sup>a</sup> | 20.3 (33.8) | 17.5 (30.0) |
| R<sub>free</sub> [%]<sup>a</sup> | 25.7 (34.0) | 21.5 (37.2) |
| **Contents of Asymmetric Unit** |  |  |
| Molecules, residues, atoms | 2; 477; 4049 | 1; 244; 2037 |
| Mg<sup>2+</sup>, GlcNAc molecules, glycerol | 2; 5; - | -; -; 4 |
| Water molecules | 132 | 174 |
| **Mean Temperature factors [Å<sup>2</sup>]<sup>b</sup>** |  |  |
| All atoms | 58.7 | 31.1 |
| Macromolecules | 58 | 30.4 |
| Ligands | 106.7 | 49.9 |
| Water oxygens | 53.5 | 36 |
| **RMSD from target geometry** |  |  |
| Bond length [Å] | 0.007 | 0.012 |
| Bond angles [°] | 0.84 | 1.04 |
| **Validation statistics<sup>c</sup>** |  |  |
| **Ramachandran plot** |  |  |
| Residues in allowed regions [%] | 2.8 | 2.5 |
| Residues in favored regions [%] | 97.2 | 97.5 |
| MOLPROBITY clashscore<sup>d</sup> | 3.23 | 3.9 |

<sup>a</sup> Data for the highest resolution shell in parenthesis
<sup>b</sup> Calculated with PHENIX [33]
<sup>c</sup> Calculated with MOLPROBITY [34]
<sup>d</sup> Clash score is the number of serious steric overlaps (> 0.4) per 1 000 atoms.

GlycoproteinD4 was crystallized with only one protein per asymmetric unit. The structure was determined at a resolution of 1.9 Å (Table 2, PDB-ID 6TM8, Figure 2 B) using the coordinates of gD1 structure for molecular replacement and refined to an R_work_ of 17.5% and R_free_ of 21.5% (Table 2). In total 244 residues could be modeled (R34 to R277). In the structures of gD1 and gD4, six cysteines were found to form three disulfide bonds at sites conserved in members of the gD plolypeptide family: C87/C209, C126/C223, and C138/C147 [27,29,32]. The overall folds of gD1 and gD4 are very similar with a root-mean-square deviation (rmsd) of 0.7 Å for 220 common Cα atoms (Figure 2 C). The cores consist of a nine-stranded (A’, B, C, C’, C”, D, E, F, and G) β-barrel, arranged in a typical V-like Ig fold, flanked by N- and C-terminal extensions with loops, α-helices (α1, α2, α3’, and α3), and small β-strands (str2-4). The termini in both structures point in opposite directions (Figure 2) and the unresolved C-termini are predicted to be unstructured by FoldIndex (https://fold.weizmann.ac.il/fldbin/findex).

### Comparison of gD1, gD4, and homolog structures

The amino acid sequence identity between EHV-1 and EHV-4 gDs is 76%, much higher than compared to HSV-1 (25%, GenBank AAK19597), PrV (34%, GenBank AEM64108) or BoHV-1 (31%, GenBank NP045370). While the global folds of gDs of these different viruses are very similar (Figure 3), the number and positions of α-helices differ. Compared to EHV-1 and EHV-4 gDs, there is an extra helix termed α1’ present in PrV and BoHV-1 gDs. PrV gD has an exclusive α2’ helix which cannot be observed in other gD structures elucidated so far. In HSV-1 and HSV-2 gDs, the α3’ helix is missing but is present in EHV-1, EHV-4, PrV, and BoHV-1 gDs (Figure 3D). In HSV-1 and HSV-2 gDs, the α1 helix is split and the α3 helix is kinked in HSV-1 (Figure 3 D) which is not seen in the other known gD structures. The six disulfide bonds C87/C209, C126/C223, and C138/C147 are conserved across EHV-1, EHV-4, PrV, HSV-1, HSV-2, and BoHV-1 gDs, while the predicted and resolved glycosylation sites in the crystal structure of gD1 and gD4 are only conserved between EHV-1 and EHV-4 (N52, N61, N297, N386) (Figure S 6). Between gD1 and gD4, also the magnesium-coordinating residues seen in the gD1 monomer-monomer interface are conserved.

**Figure 3:**
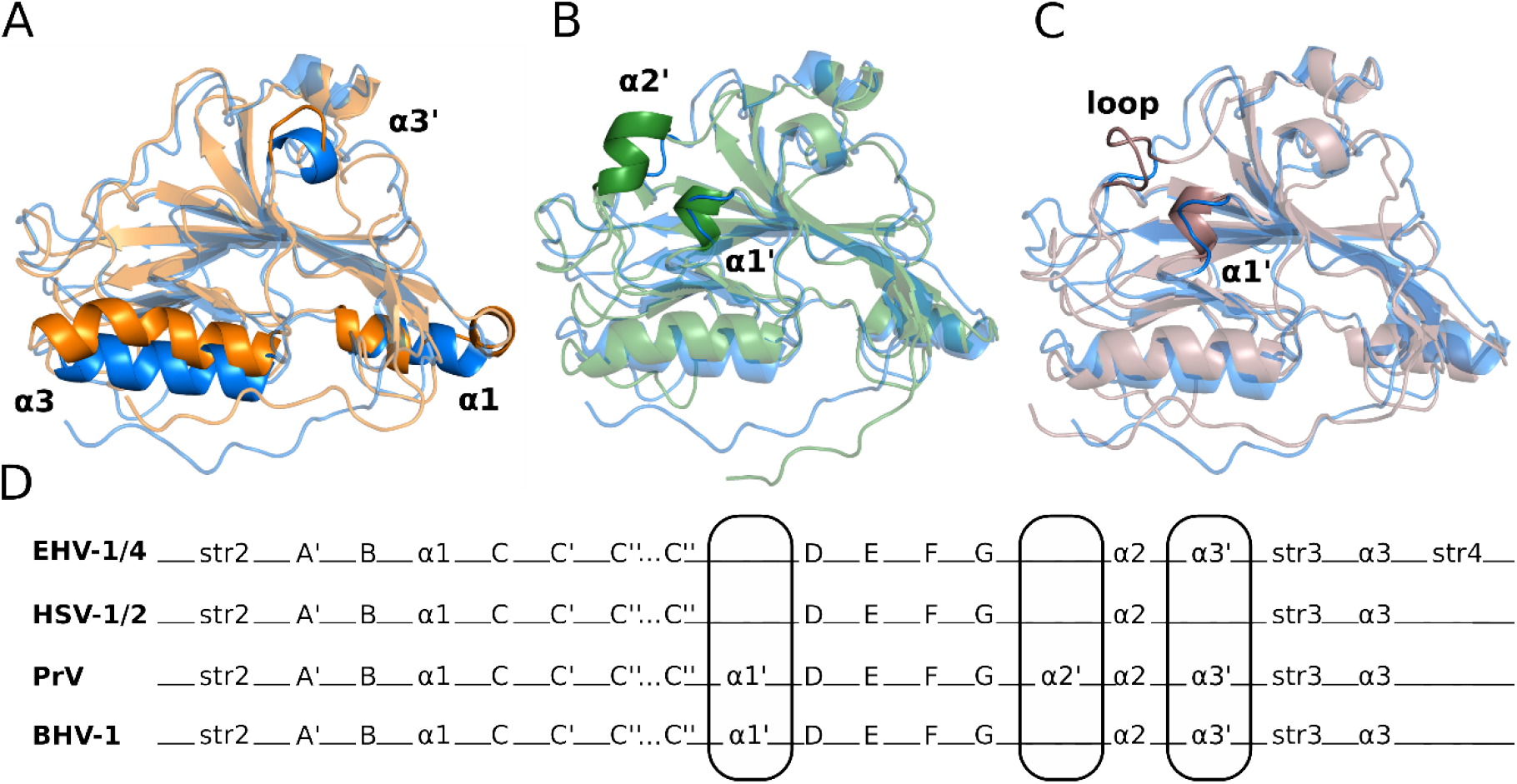
Glycoprotein D from alphaherpesviruses have a similar crystal and secondary structure. Superposition of crystal structures in cartoon representation of gD from EHV-1 (blue, PDB-ID 6SQJ) with (A) HSV-1 (orange, PDB-ID 2C36), (B) PrV (green, PDB-ID 5X5V), and (C) BoHV-1 (brown, PDB-ID 6LS9) gD. Main differences in global fold are highlighted. (D) Comparison of secondary structure elements of EHV-1/4, HSV-1/2, PrV, and BoHV-1 gD. Main differences in global fold are encircled.

### Soluble gD1 and gD4 engage recombinant MHC-I with similar binding affinities

gD-binding affinities of different alphaherpesviruses to their receptors are known to differ greatly. For example, PrV gD binds human nectin-1 in the nanomolar range [35]. HSV-1 gD interacts more weakly, in a micromolar range, with nectin-1 and herpesvirus entry mediator (HVEM) [26] similarly to BoHV-1 gD with nectin-1 [27] (Table 3). For HSV-1, HSV-2, and PrV gDs, it has been demonstrated that C-terminal truncation of the proteins increases the binding affinities up to 100-fold (Table 3).

**Table 3:**
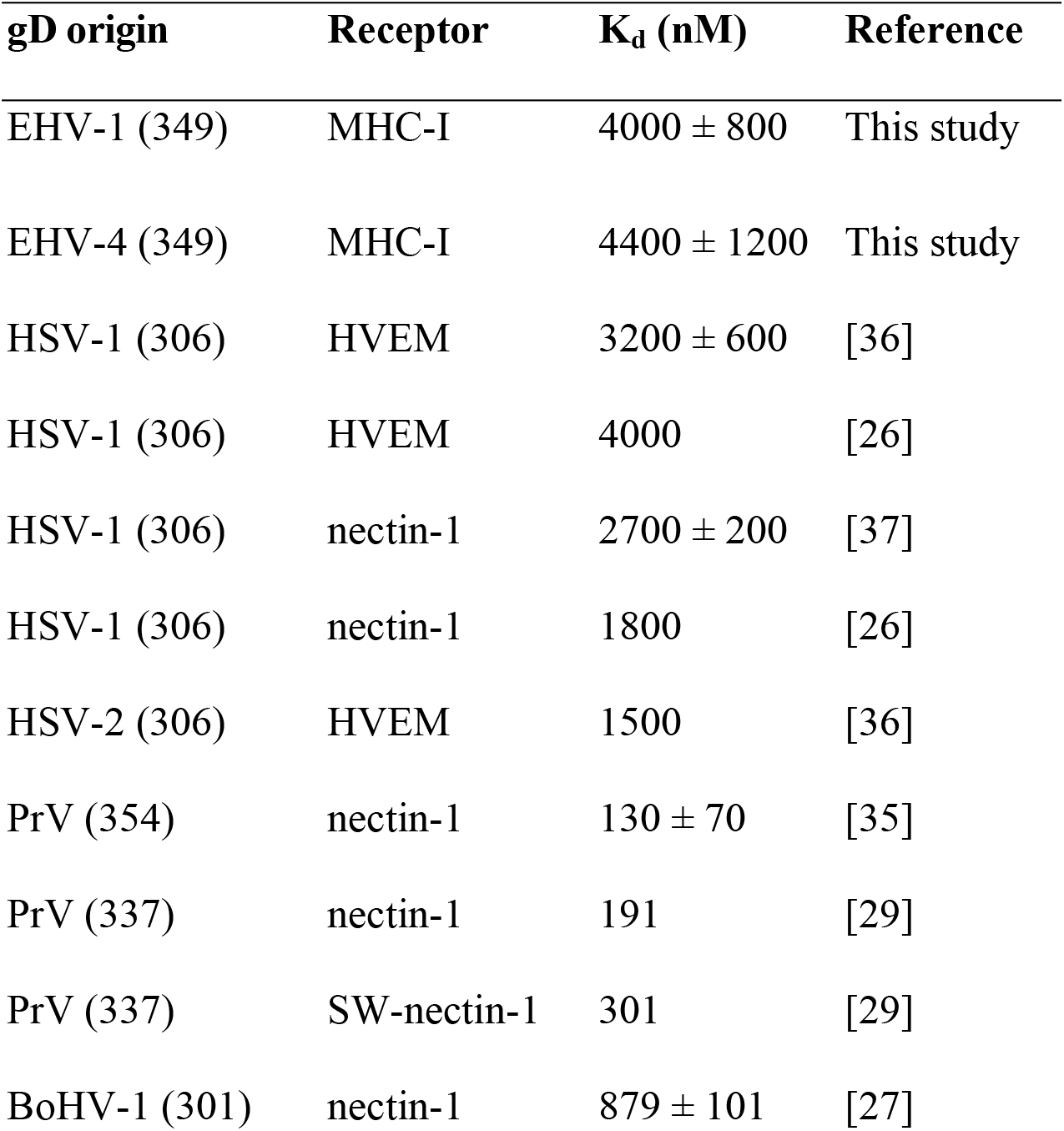
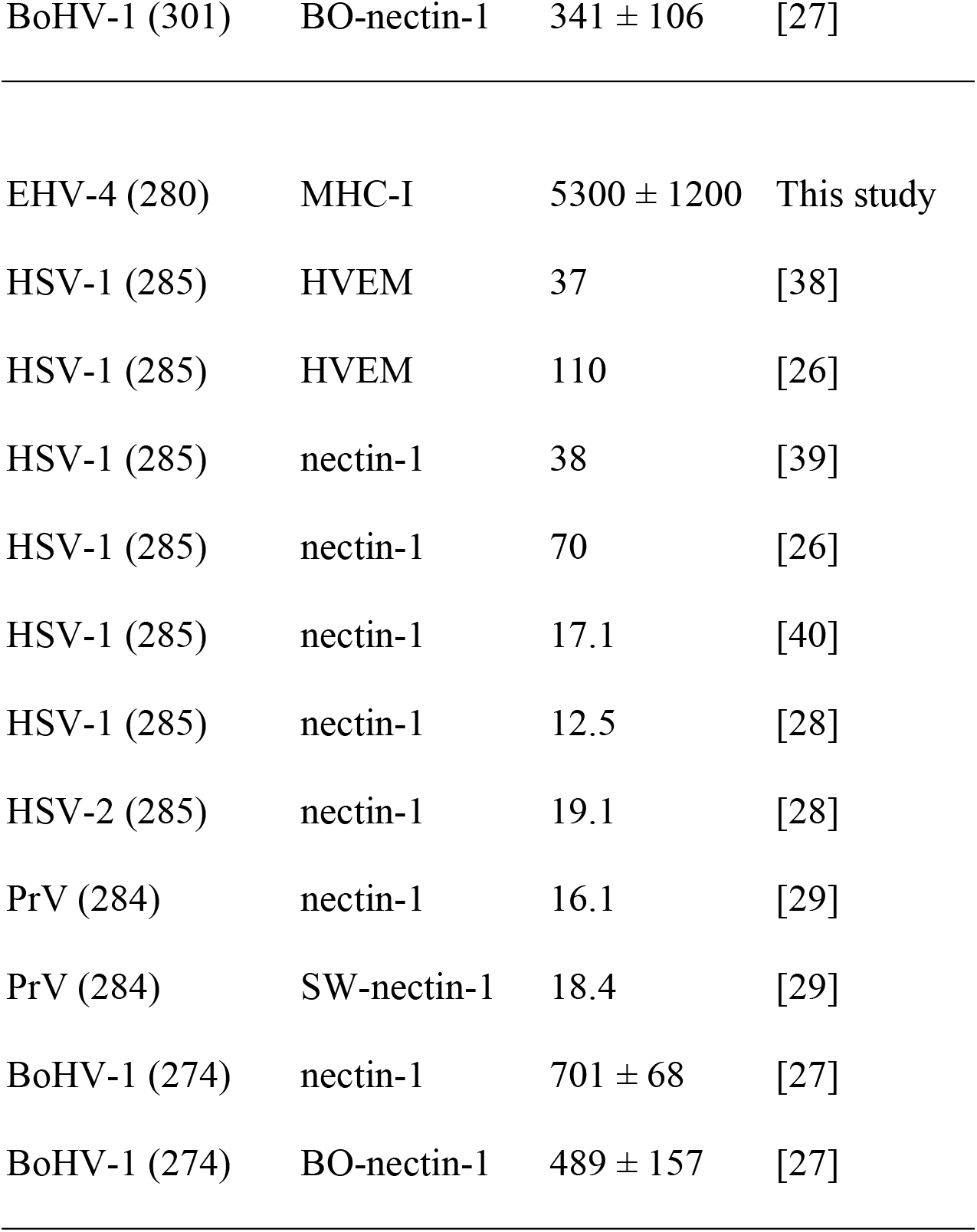
Comparison of dissociation constants (K_d_^app^) of alphaherpesviruses-gDs binding their respective receptors measured by SPR. SW = swine, BO = bovine, MHC-I = equine (Eqca-1*00101), nectin-1/HVEM = human. The C-terminal truncation of proteins is displayed in brackets under ‘gD origin’. The upper part of the table represents full-length proteins, the bottom part truncated proteins.

| **gD origin** | **Receptor** | **$K_d$ (nM)** | **Reference** |
| --- | --- | --- | --- |
| EHV-1 (349) | MHC-I | $4000 \pm 800$ | This study |
| EHV-4 (349) | MHC-I | $4400 \pm 1200$ | This study |
| HSV-1 (306) | HVEM | $3200 \pm 600$ | [36] |
| HSV-1 (306) | HVEM | 4000 | [26] |
| HSV-1 (306) | nectin-1 | $2700 \pm 200$ | [37] |
| HSV-1 (306) | nectin-1 | 1800 | [26] |
| HSV-2 (306) | HVEM | 1500 | [36] |
| PrV (354) | nectin-1 | $130 \pm 70$ | [35] |
| PrV (337) | nectin-1 | 191 | [29] |
| PrV (337) | SW-nectin-1 | 301 | [29] |
| BoHV-1 (301) | nectin-1 | $879 \pm 101$ | [27] |
| BoHV-1 (301) | BO-nectin-1 | 341 ± 106 | [27] |
| EHV-4 (280) | MHC-I | 5300 ± 1200 | This study |
| HSV-1 (285) | HVEM | 37 | [38] |
| HSV-1 (285) | HVEM | 110 | [26] |
| HSV-1 (285) | nectin-1 | 38 | [39] |
| HSV-1 (285) | nectin-1 | 70 | [26] |
| HSV-1 (285) | nectin-1 | 17.1 | [40] |
| HSV-1 (285) | nectin-1 | 12.5 | [28] |
| HSV-2 (285) | nectin-1 | 19.1 | [28] |
| PrV (284) | nectin-1 | 16.1 | [29] |
| PrV (284) | SW-nectin-1 | 18.4 | [29] |
| BoHV-1 (274) | nectin-1 | 701 ± 68 | [27] |
| BoHV-1 (274) | BO-nectin-1 | 489 ± 157 | [27] |

To study the interaction of soluble gD1 and gD4 with recombinant equine MHC-I, a surface plasmon resonance (SPR) binding assay was conducted. α-chains together with β2m with linked peptide of equine MHC-I 3.1 were produced in insect cells and purified by IMAC and SEC (Figure S 1 and Figure S 2 D). Additionally, the receptor affinity of a C-terminally truncated EHV-4 gD, gD4_36-280,_ was tested. The truncated protein was produced in the same manner as gD1 and gD4 originally with the goal to crystallize it because the flexible C-terminus was suspected to hinder crystallization of gD1 and gD4. However, shortly after the production of the truncated gD4_36-280,_ crystal structures were obtained for both gD1 and gD4 proteins. Therefore, for gD4_36-280_ only receptor binding kinetics were determined instead of crystallizing it. Another truncated version, gD4_45-276_, could not be produced in insect cells, suggesting that the protein failed to fold properly. Binding analyses for soluble gDs were conducted by using a protein dilution series in a range of 0 to 13 950 nM.

Dissociation constants (K_d_^app^) of 4000 ± 800 nM and 4400 ± 1200 nM were calculated for gD1 and gD4, respectively (Figure 4 A and B). The truncated gD4 version, gD4_36-280_, exhibited a receptor binding affinity to MHC-I in the same order of magnitude (5300 ± 1200 nM; Figure 4 C-E).

**Figure 4:**
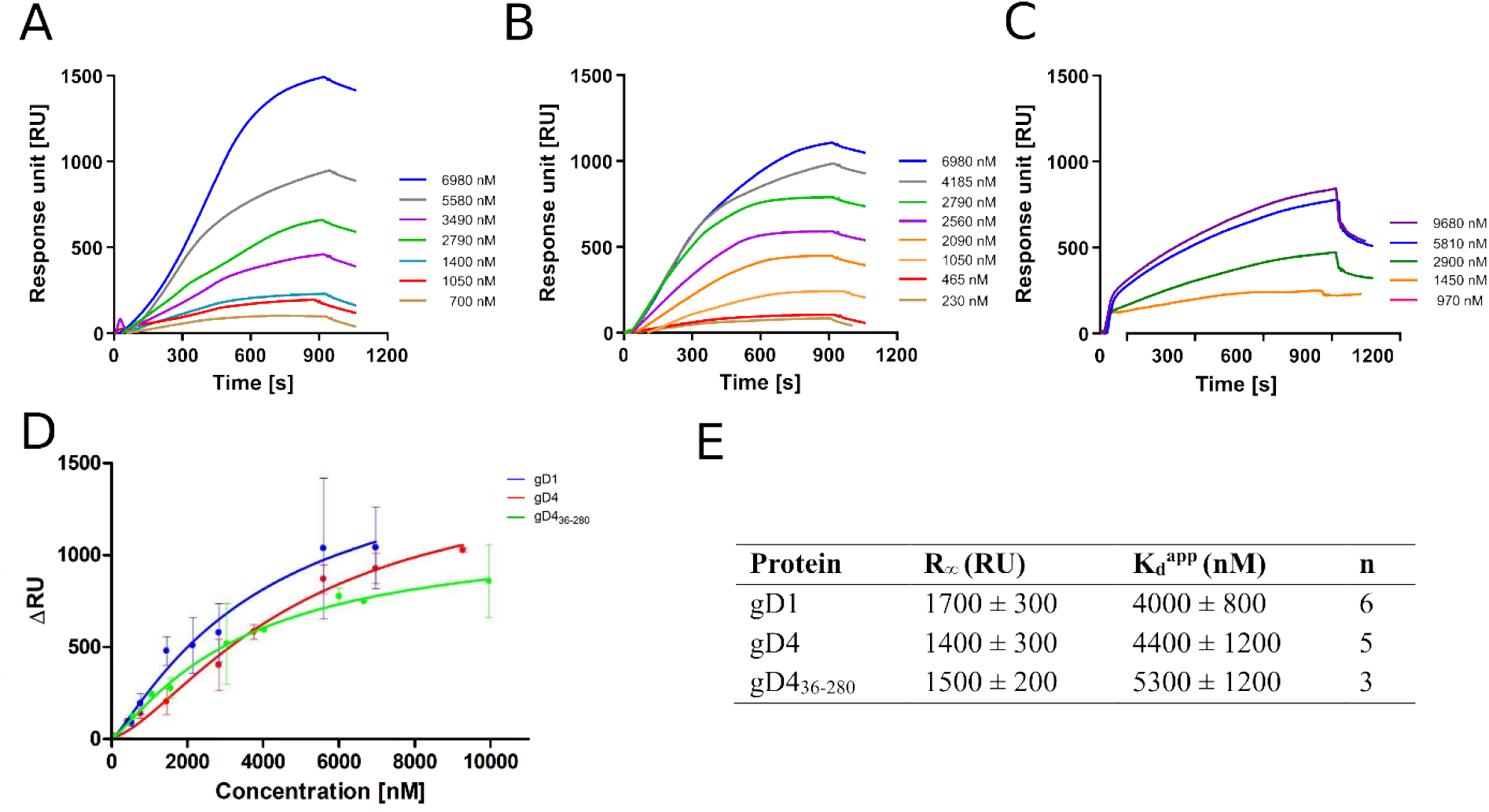
Binding affininties of gD1, gD4 and gD4_36-280_ to the entry receptor MHC-I are in micro molar range. (A, B, C) Representative SPR sensorgram profiles of recombinant gDs binding to amine-coupled recombinant MHC-I. Data were collected for several independent experiments [(A) gD1 n=6, (B) gD4 n=5, (C) gD4_36-280_ n=3). (D) Binding curves for different gD concentrations from at least three independent experiments. Displayed are means with standard deviation (SD). The solid lines represent a fit of a Hill-Wand model to the data. (E) Parameters obtained from SPR binding curves of gD1, gD4, and gD436-280. R∞ is the maximum signal obtained from the bound protein; K ^app^ is the apparent equilibrium dissociation constant, n corresponds to the number of independent experiments.

### Recombinant gD1, gD4, gD436-280 can block cell surface MHC-I

To test whether recombinant gD1 and gD4 can bind to cell surface MHC-I and subsequently inhibit virus entry, blocking assays were performed. Equine dermal (ED) cells were incubated with the recombinant proteins ranging from 0 to 150 µg/ml (0 - 3.5 µM) for one hour on ice and subsequently infected with either EHV-1 or EHV-4 at a multiplicity of infection (MOI) of 0.1. Viruses expressing green fluorescent protein (GFP) [41, 42] during early infection were used to monitor and analyze the infection levels by flow cytometry. A dose-dependent reduction of infection of up to 50% and 33% on average with 150 µg/ml protein was observed for gD1 and gD4, respectively (Figure 5 A and B).

**Figure 5:**
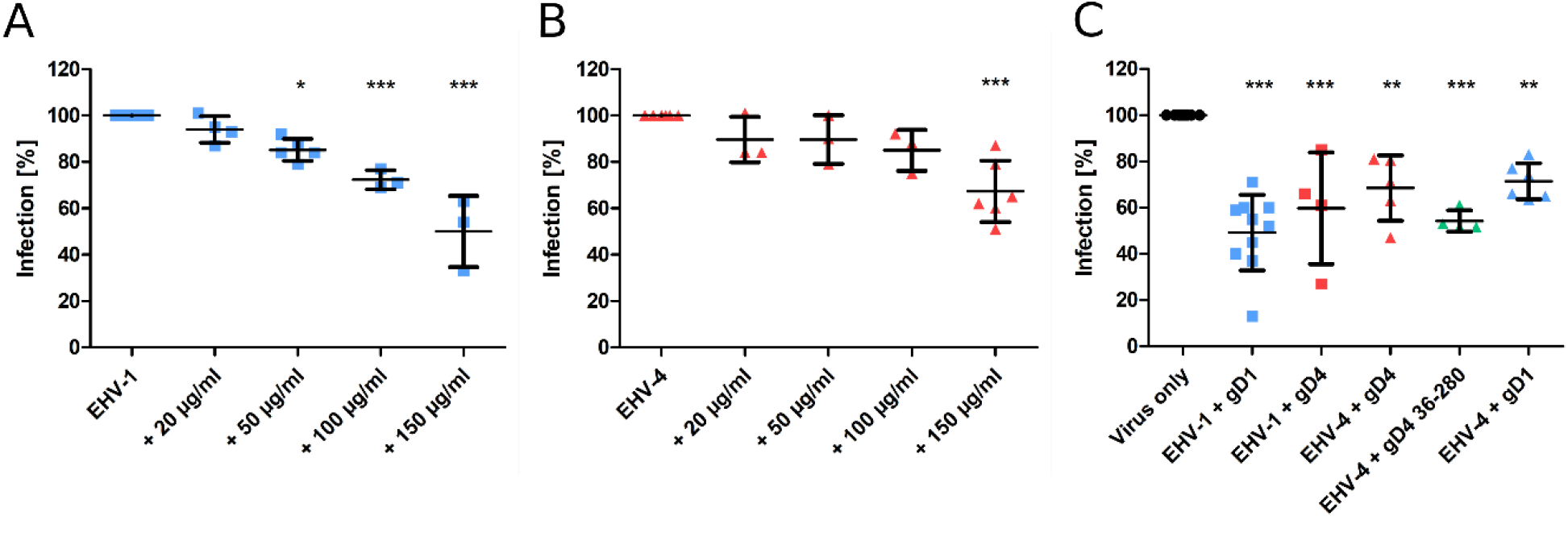
Recombinant gD1, gD4 and gD4_36-280_ blocks EHV-1 and EHV-4 infection in ED cells. (A) EHV-1 and (B) EHV-4 virus entry into ED cells blocked by different concentrations of gD1 and 4, respectively, and analyzed by flow cytometry. Cells were incubated with soluble proteins for 1 h on ice and infected with either EHV-1 or EHV-4 at MOI = 0.1. After 1 h, viruses on thecell surface were removed with citrate buffer and GFP levels were analyzed after 24-48 h by flow cytometry. (C) Plaque reduction assay of EHV-1and EHV-4 with recombinant protein. ED cells were incubated for 1 h on ice with 150 µg/ml gD1, gD4 or gD4_36-280_ and infected with 100 PFU of each virus. After 1 h, viruses on the cell surface were removed with citrate buffer and cells wereoverlaid with methylcellulose. GFP plaques were counted after 48 h. The experiment was repeated independently three times for each protein. Plaque numbers were normalized to infection levels without recombinant proteins. Statistical analysis was done using one-way ANOVA Bonferroni’s multiple comparison test, * indicates P≤0.05, ** indicates P≤0.01, *** indicates P ≤0.001. Error bars represent mean with SD.

Plaque reduction assays were performed by using a similar procedure. Here, 150 µg/ml of gD1, gD4, and gD4_36-280_ were used and cells were infected with 100 plaque forming units (PFU) of EHV-1 or EHV-4. In the presence of soluble gD1, plaque numbers were decreased on an average by 51% for EHV-1. For EHV-4, the infection was reduced by an average of 32% after blocking the cells with soluble gD4 recombinant protein. gD4 was also able to block the entry of EHV-1 by 40%. Likewise, gD1 reduced EHV-4 infection by 29%. In general, gD1 proved to be more efficient in blocking both virus infections. The gD4 variant lacking the C-terminal membrane-proximal residues, gD4_36-280_, could inhibit EHV-4 infection more efficiently with an average of 46% and proved to be slightly more potent than the extracellular domain of gD4 (32%) (Figure 5 C).

Taken together, all recombinant gDs compete with viral native proteins. A dose-dependent reduction of infection was observed for gD1 and gD4. Notably, both recombinant gDs are able to efficiently block the entry of EHV-1 and EHV-4.

### In silico modeling predicts gD1 and gD4 residues F213 and D261 as hot spots for MHC-I binding

No structures are available for gD1 or gD4 in complex with MHC-I. Therefore, protein-protein docking experiments were performed. Based on data from EHV-1 and EHV-4 mutational studies with diverse MHC-I genotypes it can be assumed that gD binds in close proximity to MHC-I A173 since genotypes with other residues at this position are highly resistant against infections [7, 18]. Available crystal structures of equine MHC-I Eqca-N*00602 (1.18.7–6) and Eqca-N*00601 (10.18) [43] feature a glutamic acid and a threonine residue at position 173 in the α2 chain, respectively, and are known not to support EHV-1 and EHV-4 infection [7]. Additionally, they contain mouse instead of equine β2m. Therefore, for *in silico* modeling of gD1 and gD4 binding MHC-I, a homology model of MHC-I genotype 3.1 was constructed to reproduce the experimental setup from the *in vitro* assays. The α-chain template (PDB ID: 4ZUU [43] and target sequences showed 85% identity and 88% similarity allowing the development of a high confidence model. In the next step, we built a homology model of equine β2m to achieve a physiological MHC-I state. The β2m template (PDB ID: 4ZUU) and target sequences showed 63% identity and 82% similarity and were therefore also highly suitable for homology modeling. The final homology models of the α-chain and β2m contained no Ramachandran outliers [44] (Figure S 7). The calculated backbone rmsd to the template amounted to 0.4 Å in each structure suggesting the correct global fold of the model. All positions (101-164 and 203-259 in MHC-I and 25-80 in β2m) and geometries of disulfide bonds considered typical for MHC were correct, suggesting a high model quality. The final model structure was obtained after assembling both chains and relaxing the homology model with a molecular dynamics (MD) simulation and used directly for the gD docking.

As described in the Materials and Methods section (*in silico* modeling suggests low impact of MHC-I peptide on gD-MHC-I binding), we initially identified gD residue D261 as a plausible hot spot contacting R169 in a peptide-free MHC-I docking. In order to mimic the experimental setup in a more realistic way, we inserted the peptide SDYVKVSNI into the MHC-I homology model for docking. This nonapeptide binds in a cleft between α2 and α3 helices of MHC-I and was used in the cell-based assays. Since the peptide conformation is strongly dependent on the peptide length [43], the peptide CTSEEMNAF from MHC-I 1.18.7–6 (PDB-ID: 4ZUU) was used to build a plausible model. The modeled peptide SDYVKVSNI showed no steric clashes and exhibited reasonable bond geometries with a negligible deviation of backbone atom positions (calculated backbone rmsd of 0.8 Å to the template) (**Error! Reference source not found.**).

gD1 and gD4 were docked to the MHC-I homology model with the modeled peptide. Subsequently, a structure showing initially identified contacts between MHC-I R169 and gD D261 were searched. Since ionic interactions are considered important for long-range binding partner recognition [45], it was assumed that this contact should be present in the protein-protein-interaction (PPI). In total, five complexes for gD1 and 21 complexes for gD4, respectively, were found to bind to MHC-I. All structures were visually inspected according to selection criteria outlined in Table 5 and one docking result was selected for each protein complex. The EHV-1 gD complex was one of the 3% best scored results and the EHV-4 gD complex was in the 11% top results suggesting that both docking solutions represent low-energy protein complexes. Both structures showed similar gD-MHC-I orientations (Figure 6 A, C) and recurring comparable contacts over the trajectories of MD simulations (**Error!** Reference source not found.).

**Figure 6:**
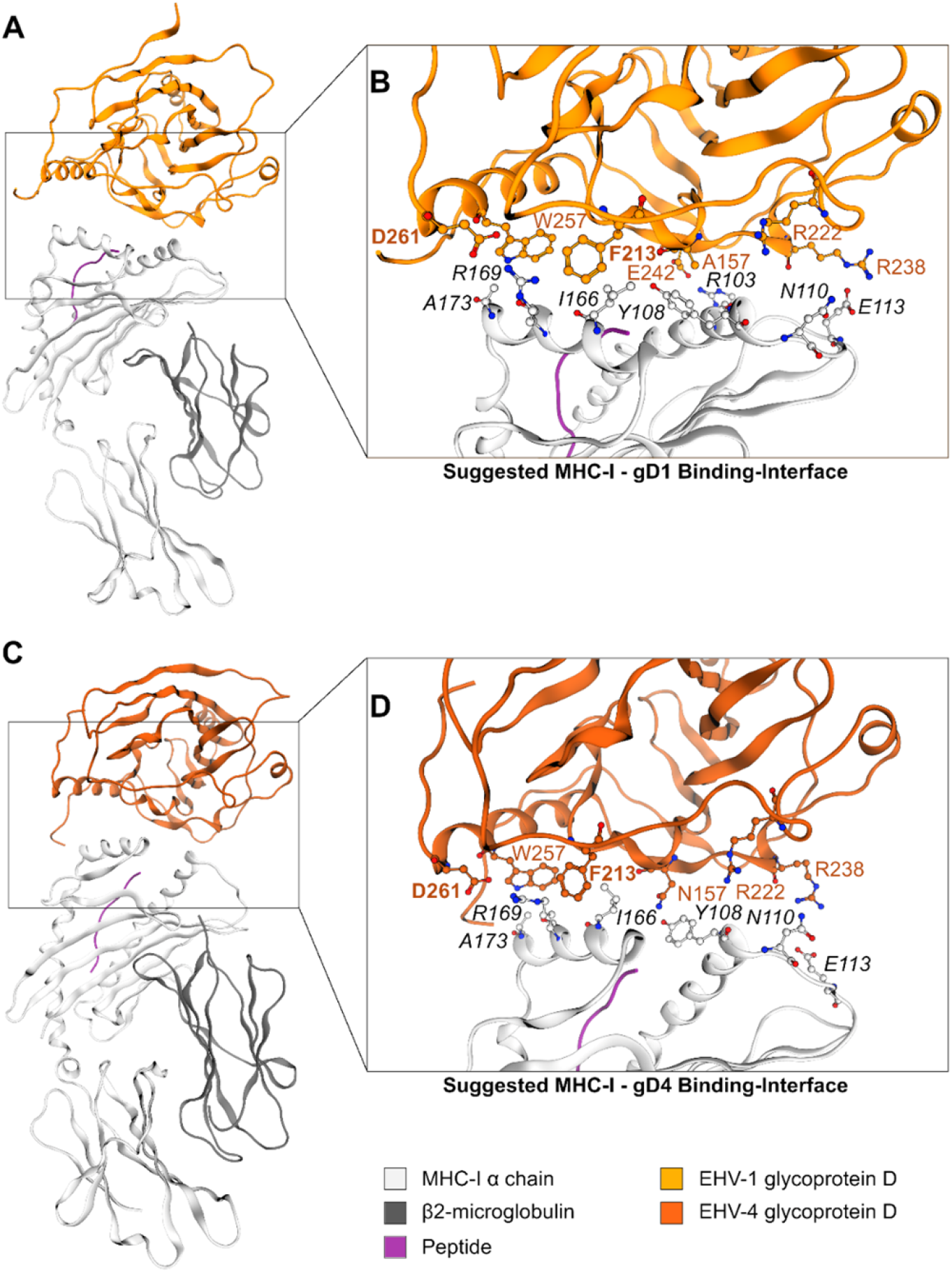
The gD1- and gD4-MHC-I interface. (A) Suggested model of the MHC-I – gD1 complex and (B) detailed view on the hypothesized binding interface. (C) Suggested model of the MHC-I – gD4 complex and (D) detailed view on the hypothesized binding interface. EHV gD residues are highlighted in orange and hot spot residues additionally in bold font. Color-code: grey ribbon – MHC-I, dark grey ribbon – β2m, orange ribbon – gD1, dark orange ribbon – gD4, purple ribbon – peptide, grey/orange balls – carbon atoms, blue balls – nitrogen atoms, red balls – oxygen atoms.

**Table 4:** Protein structures obtained from UniProt with their respective IDs.

| Protein | UniProt ID |
| --- | --- |
| MHC-I gene Eqca-1*00101 | Q30483 |
| MHC-I genotype Eqca-N*00602 | Q860N6 |
| Horse β2m | P30441 |
| Mouse β2m | P01887 |

**Table 5:**
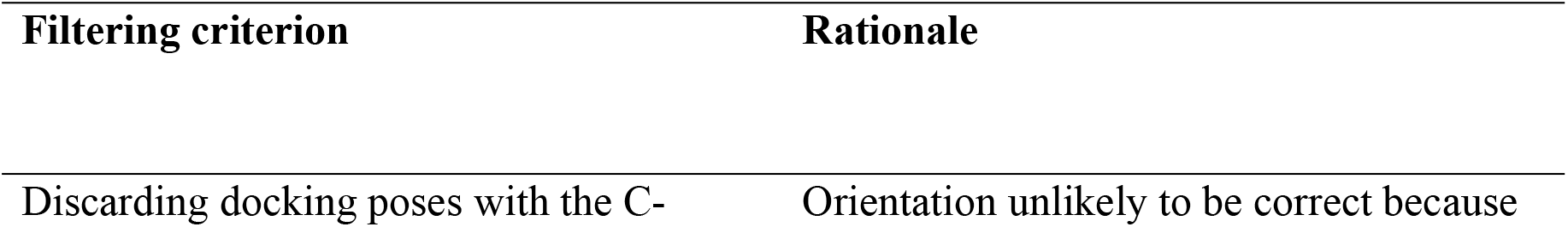
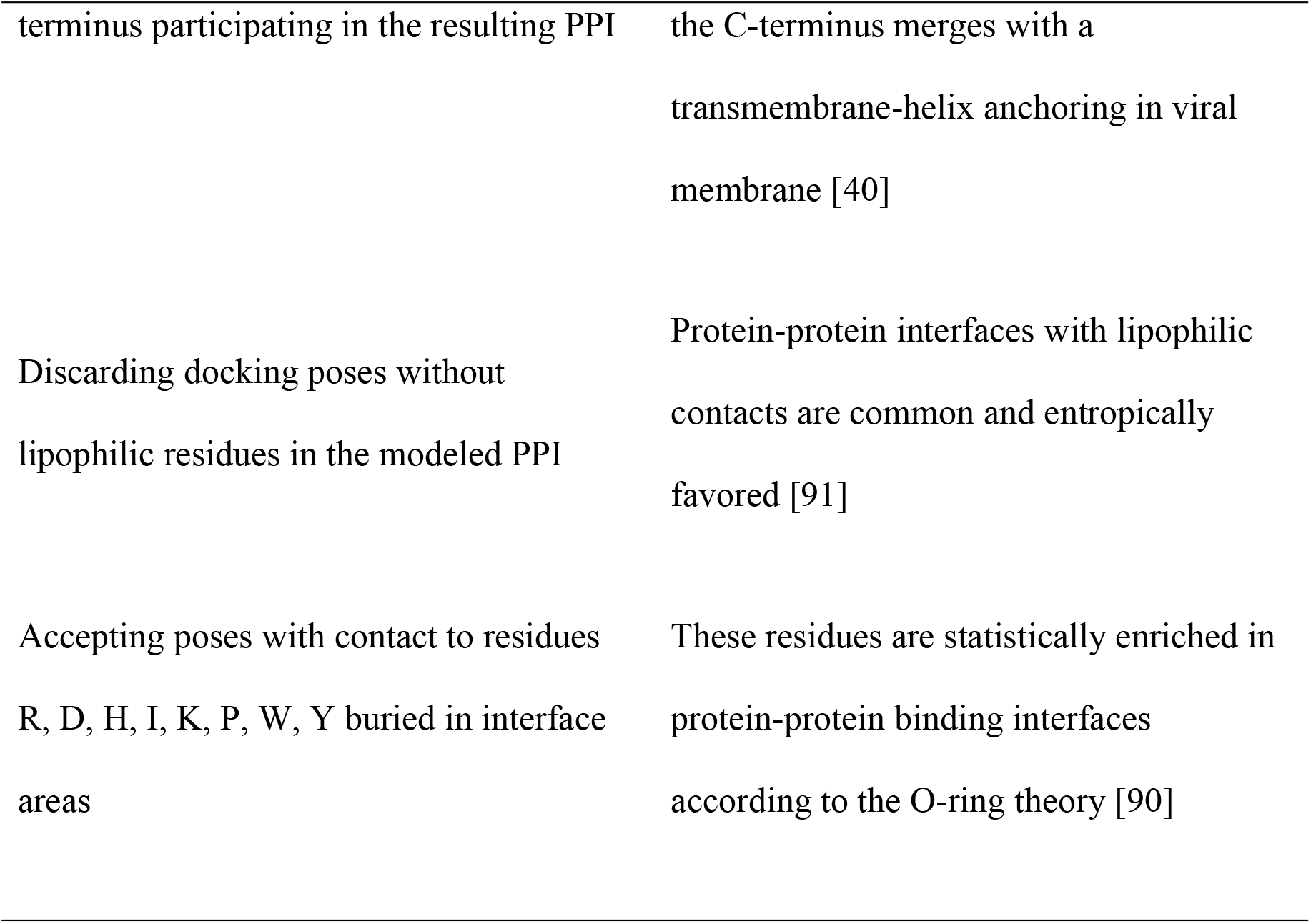
Criteria applied in our analysis of protein-protein interfaces to filter the most energetically favored docking poses.

**Table 6:** Classification of residues applied in our analysis of protein-protein interface.

| Residue type in the binding interface | Definition |
| --- | --- |
| Hot spot residues | Amino acids statistically enriched in binding sites of protein - protein complexes and contributing more than 2 kcal/mol to the binding energy [90] or form lipophilic contacts [91] |
| O-ring | Residues preventing solvation of binding hot spots [90] |
| Binding pocket | Counterpart of hot spot residues on the surface of the binding partner. |

Two contacts frequently observed between gD1 and MHC-I were identified using PyContact [46]. The first hot spot residue is D261 surrounded by the assumed O-ring including residues F213 and W257 which are contacting MHC-I binding pocket residue R169. The second hot spot residue is F213 surrounded by the assumed O-ring including Y108 and N110 and contacting MHC-I binding pocket residue I166. Additionally, an extensive hydrogen bond network between residues R103 – E242 and E113 – R238 (Figure 6, B and D) was detected (MHC-I – gD residues, respectively). All contacts and their frequencies over the trajectory of MD simulations are summarized in **Error! Reference source not found.**.

PPIs over the course of MD simulations revealed minor movements measured as backbone rmsd of maximal 3.5 Å and 6.5 Å (gD1 and gD4, respectively; **Error! Reference source not found.**C and D). Based on the optimized docking models, two gD variants were designed for experimental validation in the next step: F213A and D261N. Both residue exchanges are predicted to disrupt gD – MHC-I contacts and lead to inhibition of viral replication in a cell-based assay.

### Mutating F213A and D261N in EHV-1 and 4 gD leads to growth defects

The gD1/4-MHC-I binding hypotheses (Figure 6) were experimentally investigated by mutating the proposed key residues F213 to alanine and D261 to asparagine in EHV-1 and EHV-4 gDs. Two-step Red-mediated mutagenesis [47] was performed on EHV-1 and EHV-4 bacterial artificial chromosomes (BACs) and multi-step growth kinetics with plaque reduction assays were used for virus characterization.

All mutant viruses were successfully reconstituted from mutated BACs and the modified gD gene sequences were confirmed by Sanger sequencing. EHV-1-gD_F213A_ displayed a significant 2-log reduction in growth kinetics and low titers in cell supernatant compared to the parental virus. Reverting the mutation rescued virus growth in cell culture (Figure 7 B). Plaque sizes of wild type, mutant and revertant viruses were similar. The virus mutants EHV-1-gD_D261N_, EHV-4-gD_D261N_ and EHV-4-gD_F213A_ did not grow in cells to the extent where growth kinetics could be analyzed. However, reverting the residue exchange in EHV-1-gD_D261N_ rescued virus growth (Figure 7 A). Taken together, the gD_D261N_ and gD_F213A_ variants lead to replication-deficient viruses in EHV-1 and EHV-4.

**Figure 7:**
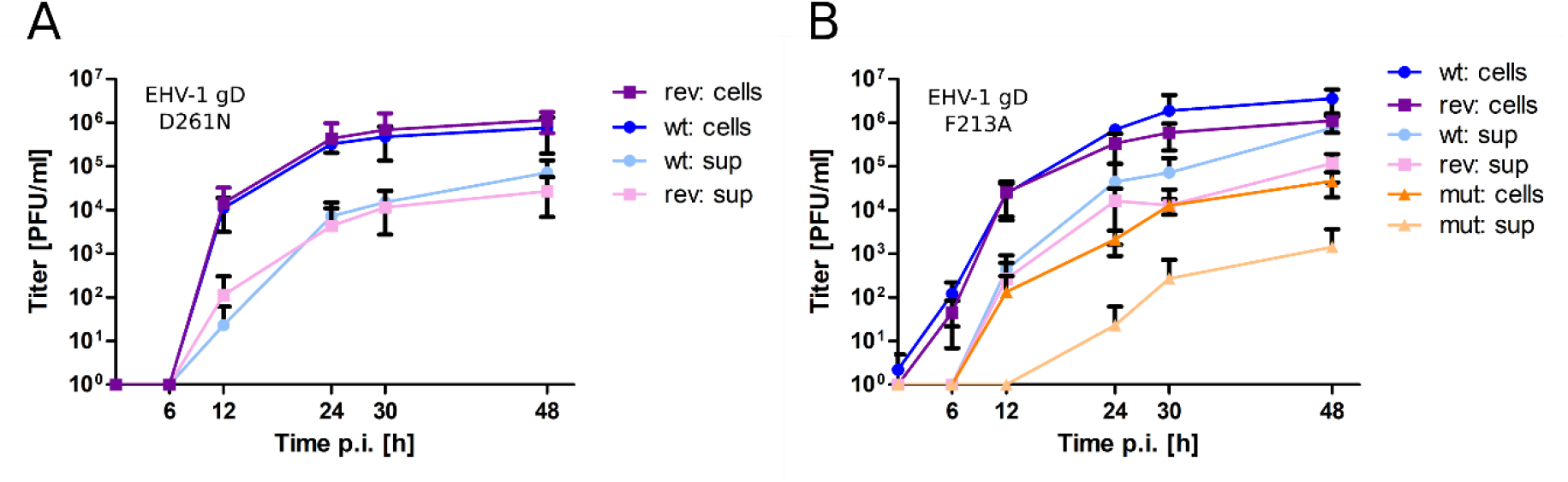
Mutating gD1 residues D261N and F213A impairs EHV-1 growth in ED cells. Multi-step growth kinetics of EHV-1 parental virus and gD mutants. ED cells were infected with an MOI of 0.01, cells and supernatant were collected separately at indicated time points post infection and titrated on ED cells. Shown are means with standard deviation (SD) of three independent experiments. (A) EHV-1 parental virus (blue colors) and EHV-1-gD_D261N_ (violet colors). (B) EHV-1 parental virus (blue colors), EHV-1-gD_F213A_ (orange colors) and EHV-1-gD-_F213A_ (violet colors).

## Discussion

Although details of cell entry of alphaherpesviruses can differ greatly between virus species, four common steps characterize the whole entry processes. First gB and/or gC attach in a relatively unspecific and reversible manner to cell surface heparan sulfate proteoglycans (HSPG) and chondroitin sulfate proteoglycans (CSPG) [48–50]. This charge-based interaction is stabilized by a stronger and specific receptor-ligand interaction [51] followed by a signaling cascade which is activated by gD and gH/gL [52]. The latter process leads ultimately to the fusion of the viral envelope with the cell membrane or in some cases to entry via endocytosis through gB [4, 53], gD is the essential protein that, in case of EHV-1 and EHV-4, binds to equine MHC-I [5–7]. The mode of gD binding to MHC-I remains elusive, although the structural understanding of alphaherpesviral gDs binding to their putative receptors has been largely extended in the last years [26,27,29,32]. Here, we present crystal structures of EHV-1 and EHV-4 gDs and propose a binding model to equine MHC-I through the key residues F213 and D261.

The crystal structures of EHV-1 and EHV-4 gDs revealed an IgV-like fold with large N- and C-termini wrapping around the core which is common for members of the gD polypeptide family. Despite high variability in sequence identities, the overall structure of alphaherpesviral gDs is conserved with only small variations in the loop regions and number of helices [26,27,29,32]. The gD termini have been shown to be important for the entry process in HSV-1 [26]. To allow receptor binding, the C-terminus needs to be displaced to free the N-terminal binding site. This could be a mechanism to prevent early onset of the fusion process before the ligand and receptor are in proximity. Subsequently, the displacement of the C-terminus allows the formation of an N-terminal hairpin loop that is crucial for HVEM binding, since exclusively gD N-terminal residues (7-15 and 24-32) interact with the receptor [26,32,54]. The displacement of the C-terminal tail is also needed for the complex formation with nectin-1 as the binding sites overlap with those of HVEM with additional C-terminal interactions (residues 35–38, 199–201, 214–217, 219–221, 223) [55]. The formation of an N-terminal loop is not involved in nectin-1 binding since the deletion of residues 7-32 had little impact on the interaction [56]. The N-terminus of gD1 and gD4 is, similarly to PrV gD, shorter than in HSV-1 gD, suggesting that HVEM cannot function as an entry receptor in these viruses. In fact, it has already been experimentally observed that HVEM is not used as entry receptors by PrV [29]. Similarly, we observed that EHV-1 also does not employ the equine HVEM homolog either [53]. In HSV-1, gD forms a dimer in the unbound state on the virus envelope [57]. This is thought to stabilize the C-terminus since viruses with a destabilized terminus could not efficiently enter cells. Although the ionic contact and high Complex Formation Significance Score of the here solved EHV-1 gD dimer interface suggest a similar function, no dimer was observed in SEC, SEC-MALS, and MS analysis. Therefore, we suppose that gD1 has no dimeric form on the virus envelope.

In contrast to results from C-terminally truncated gD homologs, which display a dramatic increase in receptor affinity (Table 3), truncated gD4_36-280_ binds MHC-I similarly as the non - truncated version. The higher affinities in the truncated homologs are explained by a faster interaction with the receptors, since the C-terminus that blocks the binding site is not required to be displaced upon binding anymore [58]. That the C-terminal truncation had no significant effect on the receptor affinity of EHV-4 gD suggests that the mode of binding differs from other alphaherpesviruses. Taking into account that EHV-1 and EHV-4 bind to MHC-I instead of HVEM or nectin-1, a different binding mechanism would be assumed. In linewith gD4_36-280_, truncated BoHV-1 gD interaction with nectin-1 showed no increased receptor affinity [27] (Table 3). A conformational change in the loop region between the G-strand and α2 helix is needed for receptor binding [27] and might explain why the affinity of the truncated BoHV-1 gD does not increase.

SPR analysis showed binding of recombinant gD1, gD4, and gD4_36-280_ to recombinant MHC-I with micromolar affinities. The K_d_ ^app^ are higher than in gD homologs of HSV-1, HSV-2, and PrV binding nectin-1. However, HSV-1 gD binding to HVEM displays similar affinities (Table 3 upper part). Nevertheless, there are two limitations of the SPR analysis in this study. First, the MHC-I molecule Eqca-1*00101 (3.1) used here allowed lower infection rates in a previous study than the molecule Eqca-16*00101 (2.16) [7]. Due to construct design reasons, the gene 3.1 fitted the purpose of crystallography better, although, no crystal structure could be obtained. However, the 2.16 molecule should display higher receptor affinities than the one observed in this study. Second, the linker region (GGGSGGGSGGGS), inserted to tether the peptide in the MHC-I complex to the β2m C-terminus, might interfere with gD receptor binding. Our attempt to model the linker to the MHC-I molecule that binds gD1 in the position hypothesized here support this hypothesis. Nevertheless, the results from blocking assays confirm that the receptor affinities of soluble gDs are unlikely to be in the nanomolar range. Furthermore, blocking assays revealed that gD1, gD4, and gD4_36-280_ can block cell surface MHC-I and thus compete with native gD in the viral envelope. It could be demonstrated that gD1 can block EHV-4 infections and vice versa implying that the receptor interaction is very similar in both viruses. This finding is supported by the binding models presented here.

The finding in SPR analysis that the C-terminally truncated gD4 does not display an increased receptor affinity was confirmed in blocking assays, thus suggesting that the receptor-binding mode differs from HSV and PrV, which is not surprising as they enter through different receptors.

The proposed docking position of gD1 to MHC-I explains why MHC-I Eqca-16*00101 (2.16) allows higher infection rates than Eqca-1*00101 (3.1) [7]. The residue 103 in the 3.1 α1 region is an arginine, which is more spacious than asparagine in 2.16, thus preventing a closer interaction with gD and leading to lower receptor binding affinities. A binding hypothesis with MHC-I 2.16 and a crystal structure of this molecule could confirm that theory. A173 of MHC-I has been shown previously to play a major role in the entry of EHV-1 and EHV-4 by two studies. First, the entry of EHV-1 into usually non-susceptible NIH3T3 cells transfected with altered hamster MHC-I Q173A has been shown together with the negative effect on infection rates of hydroyphilic residues at position 173 in equine MHC-I [18]. Second, it has been demonstrated that not all equine MHC-I genes support entry of EHV-1 and 4 into equine MHC-I transfected mouse mastocytoma (P815) cells and that MHC-I genes harboring residues other than alanine at position 173 are highly resistant against EHV-1 and 4 infections [7]. The gD1/4-MHC-I binding hypotheses explain the role of MHC-I A173 well by showing that bulkier amino acids at that position lead to steric hindrance in the gD binding pocket. This applies to MHC-I alleles 3.3 (V173), 3.4 (T173), 3.5 (E173), and 3.6 (V173), which do not support EHV-1 and 4 entry [7, 18]. The model can even explain why the genotype Eqca-7*00201 (3.7), although harboring an alanine at position 173, does not allow entry of EHV-1 and 4 into P815 3.7 [7]. The glutamine residue at position 174 is assumed to hinder gD binding sterically. The side-chain would point in the bound state into a hydrophobic residue-patch (W253, F256, W257) of gD, leading to an enthalpic penalty. Strangely, the inability of the viruses to enter via the MHC-I haplotype Eqca-2*00101 (3.2) which harbors A173 and A174 cannot be explained by the binding model. The topology of this MHC-I molecule is predicted to be very similar to those allowing virus entry. A crystal structure of the 3.2 MHC-I gene might give an explanation. Mutations in the gD binding pocket R43, W253, F256, and W257 could prove useful for a more detailed evaluation of the predicted interaction with MHC-I A173.

Another observation by Sasaki et al. [18] was that the mutation W171L in equine MHC-I impairs virus entry into NIH3T3 cells transfected with this MHC-I. Although the cell surface expression of this mutant was reduced, this is still interesting since structural data show that W171 points towards the peptide in the binding groove and should therefore not be involved directly in binding gD. The tryptophan would be able to stabilize some peptides with hydrogen bonds, whereas a leucine would not. A leucine at position 171 could therefore lead to a more loosely bound peptide with a higher flexibility, resulting in an interference via the gD-MHC-I binding. This theory would suggest that the peptide in the MHC-I binding groove itself could play a role in the receptor-ligand interaction, which could be tested by using different peptides bound to MHC-I in blocking assays and by testing mutated equine MHC-I W171L in blocking assays with soluble recombinant gDs.

Considering all these results, the question arises whether EHV-1 and EHV-4 can facilitate entry through, so far, unknown non-equine MHC-I molecules. Sasaki et al. [18] demonstrated that mutated hamster MHC-I Q173A allowed low EHV-1 infection. Unfortunately, EHV-4 has not been tested in the same manner. A computational approach could be employed to search for non-equine MHC-I molecules that are similar in the binding region that is visible in the gD1/4-MHC-I binding model and be used to select promising targets for transfection/infection assays. Experimentally, EHV-1 and EHV-4 infections could be tested in cell lines from susceptible species, e.g. bovine, rabbit, monkey, pig, cat, human [59, 60], alpacas, lamas, polar bears [61], and rhinoceros [61–63] cell lines, with and without inhibited MHC-I expression by using β2m knockdown as in Sasaki et al. [6].

Taken together, the proposed docking modes of gD1 and gD4 to MHC-I can explain several experimentally obtained results and are therefore plausible. Additionally, the docking models are supported by EHV-1 and EHV-4 viruses with mutated gD_F213A_ and gD_D261N_ that all exhibited an impaired growth. A limitation in this experiment was the difficulty of reverting EHV-4 mutations to original status to confirm that the observed effect was exclusively due to the gD_F213A_ and gD_D261N_ variants. Nevertheless, it could be shown that the gD residues F213 and D261 play a key role during entry of EHV-1 and EHV-4 providing starting points for further mutational studies possibly leading to an efficient vaccine. The results presented here might also be used to generate gD-based EHV-1 and EHV-4 inhibitors for reduction of clinical symptoms in horses and non-definite hosts.

## Materials and Methods

### Viruses

EHV-1 strain RacL11 and EHV-4 strain TH20p are maintained as bacterial artificial chromosome infections clones (BAC). The viruses have GFP under the control of the HCMV major immediate-early (IE) promoter inserted into the Mini-F sequence to easily recognize infected cells. Clones were generated as described previously in Rudolph et al. and Azab et al. [41,42,64]. The viruses were grown on equine dermal (ED) cells (CCLV-RIE 1222, Federal Research Institute for Animal Health, Greifswald, Germany) at 37 *^◦^*C under a 5% CO2 atmosphere.

### Cells

ED cells were grown in Iscove’s modified Dulbecco’s medium (IMDM; Pan, Biotech, Aidenbach, Germany) containing 20% fetal bovine serum (FBS; Biochom GmBH, Berlin, Germany), 1 mM sodium pyruvate (Pan Biotech, Aidenbach, Germany), 1% nonessential amino acids (NEAA; Biochom GmBH, Berlin, Germany), and P-S solution (100 U/mL penicillin: Panreac, AppliChem GmBH, Darmstadt, Germany; 100 µg/mL streptomycin: Alfa Aesar, Thermo Fisher Scientific, Kandel, Germany (P-S) at 37*^◦^*C under a 5% CO2 atmosphere.

Human embryonic kidney (293T, ATCC CRL-11268) cells were propagated in Dulbecco’s modified Eagle’s medium (DMEM; Biochom GmBH, Berlin, Germany), supplemented with 10% FBS and P-S. Sf9 cells (IPLB-Sf21-AE, Invitrogen, Germany) were propagated in serum free Sf-900 III medium (Gibco, Thermo Fisher Scientific, New York, USA) and High Five^™^ cells (BTI-TN-5B1-4, Invitrogen, Germany) in serum free Express Five medium (Gibco, Thermo Fisher Scientific, New York, USA) at 27*^◦^*C on orbital shaker.

### Construction of expression plasmids

Constructs were amplified from insect cell codon-optimized DNA fragments (Bio Basic Inc., Canada) for protein production in High Five insect cells. Synthetic truncated genes contained the gene of interest (gD1 residues 32-249, gD4 residues 32-249, equine MHC-I 3.1,), a C-terminal TEV protease cleavage site (ENLYFQG), and a hexa-histidine tag (His_6_), all flanked by *Eco*RI*-* and *Sca*I-restriction sites (Figure 8). Sequences of codon optimized genes can be found in the supplementary data. A further truncated form of gD4 containing the residues 36- 280 was amplified from gD4 synthetic gene with the primer pair VK50/VK56 (**Error!** Reference source not found.).

**Figure 8:**
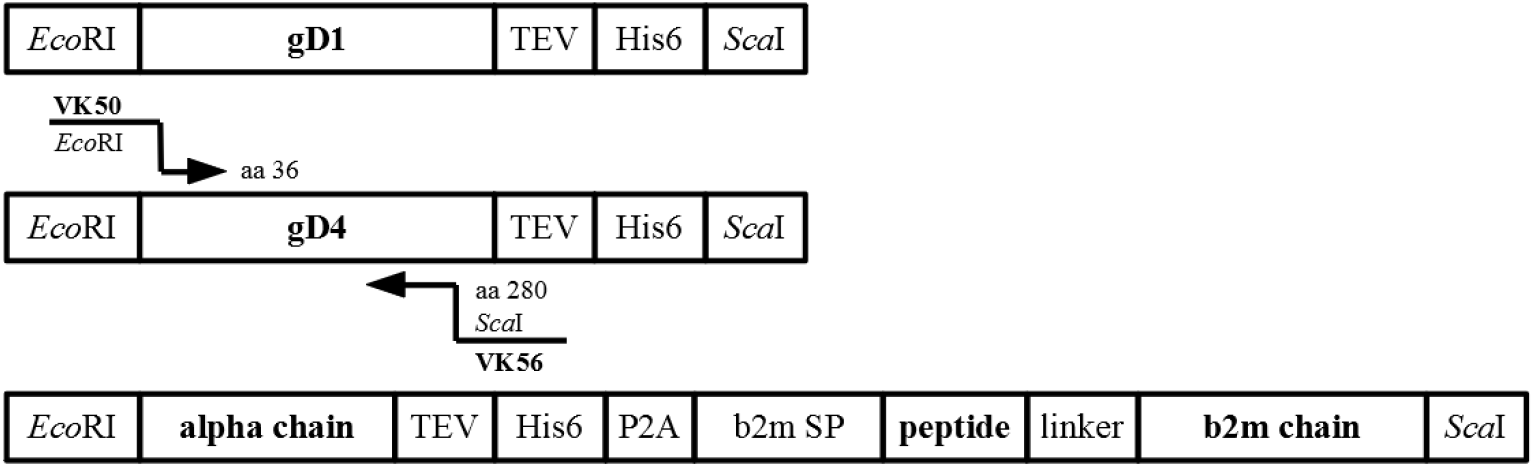
Synthetic genes used for cloning. Schematic representation of synthetic genes for protein production of gD1, gD4, equine MHC-I 3.1 with cloning strategy for gD436-280.

The Autographa californica nuclear polyhedrosis virus (AcNPV) baculovirus gp64 signal sequence under control of the very late polyhedrin promoter was inserted into the insect cell vector plasmid pACEBac1 (Addgene, LGC Standards Teddington, UK) by using another synthetic gene (VK18, LGC Genomics, Berlin, Germany) and the primer pairs VK7/VK7 (Error! Reference source not found.). Subsequently, plasmids containing synthetic genes (gD1, gD4, MHC-I) were amplified in *Escherichia coli* (E. coli), purified, and digested with *Eco*RI- and *Sca*I-restriction enzymes for insertion into the transfer vectors which was digested with the same restriction enzymes. After ligation, these plasmids were transformed into DH10MultiBac electrocompetent cells and recombinant baculoviruses produced according to manufacturer’s instructions (Bac-to-Bac expression system, Invitrogen). All constructs were verified by sequencing (VK8 or VK10/WA2, VK35/38; Error! Reference source not found.). Recombinant BACs were isolated and used for virus production in Sf9 cells as described in Santos et al. [65].

### Protein production and purification

Equine MHC-I, gD1_32-349_, gD4_32-349_, and gD4_36-280_ were expressed in HighFive cells. Cell supernatant was harvested after 72 h post infection by centrifugation. The pH was adjusted to 7 with 1M tris(hydroxymethyl)aminomethan (Tris)-HCl buffer at pH 9 on ice and incubated for at least 1 h with washed Ni^2+^-NTA beads for affinity chromatography. Beads with bound recombinant protein were collected by a gravity flow column and the proteins were eluted with a buffer containing 20 mM Tris-HCl at pH 7.5 or 2-(N-morpholino)ethanesulfonic acid (MES) at pH 6 for gDs and MHC-I, respectively, and 200 mM NaCl, 5% glycerol, and 200 mM imidazole. Concentrated protein was loaded onto a 16/600 Superdex 200 gel filtration column (GE Healthcare Piscataway, NJ). The buffer conditions were the same as in Ni^2+^-NTA affinity chromatography but with 20 mM NaCl and no imidazole. Proteins collected from size-exclusion chromatography were concentrated (Concentrators, Amicon Ultra, Millipore, Darmstadt, Germany), aliquoted and directly used for crystallization or stored at *−*80*^◦^*C.

### Crystallization, structure determination, and refinement

Crystals of EHV-1 gD were obtained by the sitting-drop vapor-diffusion method at 18°C with a reservoir solution composed of 0.1 M Tris/HCl buffer at pH 8.5, 0.2 M MgCl, and 30% (w/v) polyethylene glycol (PEG) 4000. Crystals were cryo-protected with a solution composed of 75% mother liquor and 25% (v/v) glycerol and subsequently flash-cooled in liquid nitrogen. Synchrotron diffraction data were collected at the beamline P14 at DESY (Hamburg, Germany) and at the beamline 14-2 of the MX beamline of the BESSY II (Berlin, Germany) and processed with X-ray detector software (XDS) [66]. The structure was solved by molecular replacement with PHASER [67] using the coordinates of PDB-ID 2c36 as search model for gD1 which was then used as search model for gD4. A unique solution with two molecules in the asymmetric unit for gD1 and molecule for gD4 were subjected to the program AUTOBUILD in PHENIX [33] and manually adjusted in COOT [68]. The structures were refined by maximum-likelihood restrained refinement using PHENIX [33, 69]. Model quality was evaluated with MolProbity [70] and the JCSG validation server [71]. Secondary structure elements were assigned with DSSP [72] and for displaying sequence alignments generated by ClustalOmega [73] ALSCRIPT [74] was used. Structure figures were prepared using PyMOL [75]. Coordinates and structure factors have been deposited in the PDB for gD1 with PDB-ID 6SQJ as well as for gD4 with PDB-ID 6TM8. Diffraction images have been deposited at proteindiffraction.org (gD1: DOI 10.18430/m36sqj and gD4 DOI 10.18430/m36tm8).

### Mass spectrometry analysis

Intact protein mass of gD1, gD4, and MHC-I was determined by matrix-assisted laser desorption ionization-time of flight mass spectrometry (MALDI-TOF-MS) using an Ultraflex-II TOF/TOF instrument (Bruker Daltonics, Bremen, Germany) equipped with a 200 Hz solidstate Smart beam^™^ laser. Samples were spotted using the dried-droplet technique on sinapinic acid (SA) or 2,5-dihydroxybenzoic acid (DHB) matrix (saturated solution in 33% acetonitrile / 0,1% trifluoroacetic acid). The mass spectrometer was operated in the positive linear mode, and spectra were acquired over an m/z range of 3,000- 60,000. Data was analyzed using FlexAnalysis 2.4. software provided with the instrument. Protein identity was determined by tandem mass spectrometry (MS/MS) of in-gel digested Coomassie stained protein with 12,5µg/ml Glu-C and trypsin, and 10µg*_/_*ml Asp-N in 25nm ammonium bicarbonate.

N-terminal c and C-terminal (z+2) sequence ion series were generated by in-source decay (ISD) with 1,5-diaminonaphthalene (1,5-DAN) as matrix (20 mg/ml 1,5-DAN in 50% acetonitrile / 0,1% trifluoroacetic acid). Spectra were recorded in the positive reflector mode (RP PepMix) in the mass range 800–4,000.

### SEC-MALS analysis

For molecular mass determination of soluble, recombinant gD1, SEC-MALS [76] was performed. Protein solution was run at room temperature on a Superdex 75 10/300 GL (GE Healthcare, Piscataway, NJ) column with 2 mg*/*ml gD1 and a mobile phase composed of Tris-HCl at pH 7.5, 200 mM NaCl, 5% glycerol, and 0.02% sodium azide, attached to a high-performance liquid chromatography (HPLC) system (Agilent Technologies, Santa Barbara, CA, USA) with a mini DAWN TREOS detector (Wyatt Technology Corp., Santa Barbara, CA, USA). Data was acquired and analyzed with the ASTRA for Windows software package (version 6.1.2).

### Surface plasmon resonance

Binding kinetics of soluble gD1, gD4, and gD4_36-280_ binding to amine-coupled recombinant, equine MHC-I 3.1 were measured at 25^°^C on a surface plasmon resonance (SPR) GE Biacore J Biomolecular Interaction Analyser instrument (Uppsala, Sweden) using a polycarboxylate hydrogel sensor chip HC200M (XanTec bioanalytics GmbH). The second channel was coated with poly-L-lysine and positive nanogels (size 214nm) [77] that were shown to interact only weakly with gDs and used as negative control. The control sensorgrams were subtracted from reaction sensorgrams and normalized. The surfaces were regenerated with buffer containing 200 mM NaCl and 10 mM NaOH after each cycle. Serial dilutions of gDs ranging from 0 to 10000 nM were injected at medium flow and the interaction with MHC-I was monitored for 15 min. The response curves of gDs binding to the MHC-I were fitted to the Hill-Wand binding model 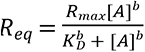 [78] using Sigma plot 12.0 software.

### Generation and analysis of gD1/4-MHC-I binding model

#### Protein data

Sequences of MHC-I and β2m were obtained from UniProt-Databank [79]. The protein sequences with their respective UniProt IDs are listed in Table 4.

#### Homology modeling

Homology models were prepared using MOE (version 2018.0101; Molecular Operating Environment, Chemical Computing Group ULC, Montreal, Canada). The models were constructed using GB/VI scoring [80] with a maximum of ten main chain models. To check geometry of obtained homology models, Ramachandran (phi-psi angle) plots [44] were calculated with MOE.

The full MHC-I gene 3.1 model was prepared based on the hybrid equine α-chain - mouse β2m X-ray crystal structure with the best resolution (PDB-ID: 4ZUU [43]). The α-chain and β2m homology models were superposed onto the template. All side chain clashes were removed by energy minimization using the OPLS-AA force field [81], resulting in the full MHC-I gene 3.1 model. The complete model was relaxed in a MD simulation on settings described below.

#### Protein-Protein Docking

MHC-I-gD1 complex were prepared with MOE 2018 by protonation [82], modeling of missing side chains, deleting water molecules and charging termini. Protein - protein docking was performed using Rosetta 3 suite (version 2018-33) [83, 84]. The orientations of MHC-I and gD were randomized (flags -randomize 1 -randomize 2) and spun (flag -spin) to the beginning of the docking process. Docking perturbation parameters were set to default: 3 Å translational and 8° rotational movement (flag -dock_pert 3 8) [85]. The residue side chains of both docking partners were allowed to rotate around the *χ*1 and the *χ*2 angles (flags -ex1 - ex2). In total 10 000 docking runs were conducted (flag -nstruct 10 000) as recommended by the Rosetta documentation [86], yielding over 7 000 poses in each docking round. A flat harmonic distance constraint between the Cα of MHC-I A173 and the gD backbone was added based on reported genotype studies indicating the pivotal role of MHC-I A173 [7, 18]. This allowed us to limit the number of possible protein-protein docking complexes and perform local docking as recommended by the Rosetta documentation [86]. Constraint parameters were set to the default [86]: Distance 0 Å, standard deviation 1 Å and tolerance 5 Å to achieve the closest possible proximity between chains. In order to obtain a full MHC peptide complex, peptide SDYVKVSNI, as used in cell-based assays, was manually fitted into the MHC-I cleft. To fit our sequence, the peptide structure from the template (PDB-ID: 4ZUU) containing a nonapeptide CTSEEMNAF was superposed on the MHC-I homology model. The co-crystallized nonapeptide sequence was manually mutated. The side chain conformations were adjusted using MOE’s rotamer tool and energy minimized using the OPLS-AA force field to relax atomic clashes.

Finally, gD1 and gD4 were docked into the prepared MHC-peptide complex. In order to find the final and most plausible docking pose of gD1 and gD4 in complex with MHC-I-peptide, an in-house developed MD Analysis-based (version 0.19.2) [87, 88] script was used to find ionic key-contact defined as a distance of maximal 4.5 Å between C*γ* atom of D261 in gD and C*ζ* atom of R169 in MHC-I. The script was run in a Python 3.6 environment [89].

#### Filtering of Docking Poses and Classification of Residues Involved in the Protein-Protein Interface

In order to filter the most plausible from all best-scored docking poses, we applied three rules based on reported statistical evaluation of various protein-protein interactions (PPIs) [90, 91] and biological function of Herpesvirus gD (Table 5) [40]. The residues involved in the protein-protein binding in the obtained binding hypotheses were classified according to the O-ring theory (Figure 6) [90].

#### Molecular Dynamics Simulations and Protein-Protein Interaction Analysis

Molecular Dynamics (MD) simulations were prepared using Maestro (version 11.7; Schrödinger, New York, USA) and carried out using Desmond 2018-3 (version 5.5) [92]. All systems were simulated on water-cooled GeForce RTX 2080 Ti graphics processing units (NVIDIA Corporation, Santa Clara, USA). The full MHC-I gene 3.1 homology model was solvated in a cubic box with 12 Å buffering with SPC water model [93]. The system was neutralized using sodium or chloride ions and osmotic pressure was adjusted with 0.15 M sodium chloride to achieve an isotonic system. The subsequent system relaxation was performed according to the default Desmond protocol. The MD simulation ran under periodic boundary conditions and as an NPT ensemble (constant particle number, pressure and temperature) using the OPLS 2005 force field [94]. The MD simulation was performed in one replicate over 100 ns. Coordinates of the relaxed model were retrieved after the backbone rmsd (**Error! Reference source not found.**A) had reached a stable plateau around 3 Å indicating protein equilibration.

Docking poses were simulated under the same conditions as the homology models. The movement of protein - protein complex hypotheses was observed in a single MD simulation over 100 ns resulting in ca. 5000 complex conformations. MD simulations of the final selected docking pose were performed in triplicates. The simulated systems contained around 140 000 – 168 500 atoms. The proteins were wrapped, aligned on the backbone and visually inspected in VMD [95] (version 1.9.3). Protein-protein interactions were analyzed using PyContact [46] (version 1.0.1) on default settings (distance cutoff 5.0 Å, angle cutoff 120.0° and distance cutoff between hydrogen and hydrogen bond acceptor of 2.5 Å). The PyContact analysis was run in a Python 2.7 environment [89].

#### *In silico* modeling suggests low impact of MHC-I peptide on gD-MHC-I binding

To test whether additional bias emerging from peptide modeling influenced docking experiments, two docking rounds were performed. First, gD1 was docked to peptide-free MHC-I. Second, gD1 was docked to MHC-I 3.1 homology model containing the peptide SDYVKVSNI to check if docking provides comparable PPIs. Both docking rounds were performed using the settings described above. As the initial filtering step, the ten highest scored docking poses with the lowest Rosetta Energy were selected [96]. For further filtering, the rules described in the Methods section were applied (Table 5). Four out of ten docking poses fulfilled all three rules. Subsequently, single MD simulations for each docking pose were performed to examine PPI stability. For all protein - protein complexes the backbone rmsd was calculated to obtain an overview of the amplitude of protein movements. Only one docking pose showed a nearly constant backbone rmsd value of 6 Å indicating low complex movement (**Error! Reference source not found.** B). In order to characterize the obtained PPI, we applied selection criteria and identified three residue patch-classes in the binding surface as described in the Methods section (Figure 6). An inspection of the PPI over an MD simulation trajectory with PyContact [46] (**Error! Reference source not found.**) revealed two gD hot spot residues: D261 (surrounded by assumed gD O-ring T161, F213, and W257 and contacting binding pocket MHC-I residue R169 over whole simulation time) and W257 (surrounded by assumed O-ring R43, T161, and F213 and contacting binding pocket MHC-I residue I166 over the whole trajectory). It can be concluded that PPIs of peptide-free and peptide-bound docking poses are formed with similar residue patches (**Error! Reference source not found.**, **Error! Reference source not found.**) suggesting that the presence of the peptide in MHC-I does not influence gD binding. The key residue F213 is involved in both PPIs indicating its importance. We observed that peptide-bound docking poses exploit larger PPIs with more possible interactions than the peptide-free docking pose. We assume that more contacts between gD and MHC-I are favorable for the binding. Therefore, peptide-bound docking poses were chosen as the final ones.

### BAC mutagenesis

The point mutations F213A and D261N in EHV-1 and EHV-4 gDs were introduced via a two-step Red recombination [47]. In brief, polymerase chain reaction (PCR) primers (**Error! Reference source not found.**) were designed in a way that the 50 nucleotide recombinantion arms include the point mutation and sequence to amplify the *kan*^R^ gene. For construction of EHV-1-gD_D261N_, EHV-1gD_F213A_, EHV-4-gD_D261N_, and EHV-4-gD_F213A_ the primer pairs VK61/VK62, VK63/VK64, VK65/VK66, and VK67/VK68 were used for PCR amplification respectively. After Dpn-1 digest of PCR products, fragments were electroporated into GS1783 containing EHV-1 or EHV-4 BACs. DNA from Kanamycin resistant colonies was extracted and correct mutants were selected based on Restriction fragment length polymorphism (RFLP) using the restriction enzyme Pst-I. Correct clones were subjected to another round of Red recombination to remove the *kan*^R^ gene. Final clones were further analyzed by RFLP and sequencing, BAC extracted, puroified and transfected into 293T cells. Cells and supernatant were harvested three days post transfection and used to infect ED cells. Revertants were produced from mutant clones using the same procedure with primer pairs VK69/VK70, VK71/VK72, VK73/VK74, and VK75/VK76 for producing EHV-1R-gD_D261N_, EHV-1R-gD_F213A_, EHV-4R-gD_D261N_, and EHV-4-RgD_F213A_, respectively. All genotypes were confirmed by PCR, RFLP, and Sanger sequencing using the primer pair WA2/VK8 and WA2/VK10 (**Error! Reference source not found.**) for EHV-1 and EHV-4 mutants, respectively.

### Western blotting

Western blot analysis was performed with soluble proteins: 50 µg*/*ml MHC-I, 5 µg*/*ml gD1, and 5 µg*/*ml gD4. Proteins were separated by 12% SDS-PAGE, transferred to a polyvinylidene difluoride (PVDF) membrane (Roth, Karlsruhe, Germany), detected with 1:1 000 dilution rabbit anti-His_6_ (Sigma-Aldrich, St Louis, USA) antibody and 1:10 000 dilution goat anti-rabbit-HRP antibody (Sigma-Aldrich, St Louis, USA) and visualized by enhanced chemiluminescence (ECL Plus; Amersham).

### Virus blocking assays

To block cell surface MHC-I, 1,5x10^5^ ED cells were seeded in 24-well plates. In the next day, cells were incubated with 20, 50, 100 or 150 µg/ml recombinant gD1, gD4 or gD4_36-280_ for 1 h on ice. Subsequently cells were infected with either EHV-1 or EHV-4 at MOI=0.1 and incubated for 1 h at 37^°^C. To remove un-penetrated viruses, cells were washed with citrate buffer, pH 3, containing 40 mM citric acid, 10 mM potassium chloride and 135 mM sodium chloride, then washed twice with phosphate buffered saline (PBS) and infection allowed to proceed for 24 h (EHV-1) or 48 h (EHV-4). For measurement of fluorescence intensity 10 000 cells were analyzed with a FACSCalibur flow cytometer (BD Biosciences) and the software CytExpert (Beckman Coulter, Krefeld). The experiment was repeated three independent times for each protein.

For plaque reduction assay the same protocol was applied for blocking surface MHC-I with minor changes. Cells were initially incubated with 150 µg*/*ml gD1 or gD4, infected with 100 PFU, and overlaid with 1.5% methylcellulose (Sigma-Aldrich, Taufkirchen, Germany) in Iscove’s Modified Dulbecco’s Medium (IMDM) after citrate treatment and washes with PBS. GFP plaques were counted after 48 h with a Zeiss Axiovert.A1 fluorescent microscope (Carl Zeiss AG, Jena, Germany). The experiment was repeated three independent times for each protein.

### Virus growth kinetics

Virus replication was tested using multi-step growth kinetics and plaque areas were obtained as described before [50]. ED cells were grown to confluency in 24-well plates, infected with an MOI of 0.1 virus and incubated for 1 h at 37^°^C. Viruses on the cell surface were removed by washing with citrate buffer. After neutralization with IMDM, cells were washed twice with PBS and finally overlaid with 500 µl IMDM. At indicated times after the citrate treatment cells and supernatant were collected separately for EHV-1 and together for EHV-4 and stored at -80*^◦^*C. Titers were determined by plating dilution series onto ED cells and counting plaque numbers after two days under a methylcellulose overlay. All plates were fixed for 10 min with 4% paraformaldehyde, washed with PBS and stained for 10 min with 0.1% crystal violet solution in PBS which was washed away with tab water. Viral titers are expressed as PFU per milliliter from three independent and blinded experiments.

### Statistical analysis

For blocking assays, plaque numbers were normalized to infection levels without recombinant proteins. Statistical analysis was done using GraphPad Prism 5 software (San Diego, CA, USA) and one-way ANOVA Bonferroni’s multiple comparison test, * indicates P*≤*0.05, ** indicates P*≤*0.01, *** indicates P*≤*0.001. Statistical analysis was done using an unpaired, one-tailed test. P<0.05 was considered significant.

## Acknowledgments

We acknowledge DESY (Hamburg, Germany), a member of the Helmholtz Association HGF, for the provision of experimental facilities. Diffraction data were collected at beamline P11 of the PETRA III storage ring (Hamburg, Germany) under proposal I-20180662, assisted by Eva Crosas, and at the beamlines of the BESSY II storage ring (Berlin, Germany) via the Joint Berlin MX-Laboratory sponsored by Helmholtz Zentrum Berlin für Materialien und Energie, Freie Universität Berlin, Humboldt-Universität zu Berlin, Max-Delbrück-Centrum für Molekulare Medizin, Leibniz-Forschungsinstitut für Molekulare Pharmakologie, Charité - Universitätmedizin Berlin and Max-Planck-Institut für Kolloid- und Grenzflächenforschung. We acknowledge David Machalz for providing us the MDAnalysis script allowing us fast docking poses analysis and Jong-Heng Huang for performing MALS experiments. For mass spectrometry (C.W.), we would like to acknowledge the assistance of the Core Facility BioSupraMol.

## Supporting information

**Figure S 1: SDS-PAGE and western blot of MHC-I, gD1, and gD4.** (A) Coomassie stained SDS-PAGE on 12% gel. For MHC-I, only the α-chain is visible in (A) and (B). M = marker. (B) Western blot: MHC-I (50 µg/ml), gD1 (5 µg/ml), and gD4 (5 µg/ml) detected with 1:1000 rabbit anti-His_6_ antibody and 1:10000 goat anti-rabbit-HRP as secondary antibody.

**Figure S 2: Protein purification by size exclusion chromatography (SEC).** Representative SEC curves of concentrated (A) gD1, (B) gD4, (C) gD4_36-280_, and (D) MHC-I run on Superdex 200 16/600 after purification through immobilized metal ion affinity chromatography (IMAC). Solid curves show UV absorbance at 280nm and the dotted curves at 260nm.

**Figure S 3: SDS-PAGE of protein purification by size exclusion chromatography (SEC).** Representative size exclusion chromatography (SEC) fractions of proteins produced in insect cells on Coomassie stained 12% sodium dodecyl sulfate (SDS) gels. (A) gD1 (approximately 43 kDa), (B) gD4 (approximately 43 kDa), (C) gD4_36-280_ (approximately 30 kDa), and (D) MHC-I (comprised of α-chain with an approximate size of 38 kDa and β2m (with the linker and peptide) with an approximate size of 13 kDa). M = marker, FT = flow through affinity chromatography, E = elution affinity chromatography, L = loaded on SEC column.

**Figure S 4: Tandem mass spectrometry (MS/MS) of in-gel digested gD1 and gD4.** Exemplary spectra are shown which confirm the identity of the analyzed proteins and the integrity of their termini. The inserts display the theoretical b and y fragments ions. (A) N-terminal peptide of gD1 generated by cleavage with Asp-N endoproteinase (M+H= 1574.90, pos. 001-013, sequence EFEKAKRAVRGRQ.D); (B) N-terminal peptide of gD4 obtained by trypsin cleavage (M+H=1013.52, pos. 001-007, sequence EFENYRR); (C) C-terminal peptide of gD4 including the His_6_-tag, generated by Glu-C endoproteinase (M+H=1563.74, pos. 323-334, sequence ENLYFQG-H_6_).

**Figure S 5: *S*ize exclusion chromatography (SEC) combined with multi-angle static light scattering (MALS) analysis of gD1.**

The gD1 crystal structure consists of a homodimer with two ions interpreted as magnesium originating from the crystallization solution, trapped between them. The ionic interaction together with a high Complex Formation Significance Score of 0.765 (PDB Proteins, Interfaces, Structures and Assemblies (PISA) server www.ebi.ac.uk/pdbe/pisa/) suggested that gD1 might form a dimer on the virus envelope as has been proposed for HSV-1 gD [26]. To evaluate whether recombinant gD of EHV-1 has a homodimeric and/or monomeric form in solution, molecular mass calculation based on SEC*-*MALS analysis was performed for gD1. Green curve represents the normalized refractive index trace (intensity, right y-axis) for gD1 eluted from a Superdex 200 10/300 column. Blue line under the peak corresponds to the averaged molecular mass distribution (left y axis) across the peak. Exclusively the monomeric form with an approximate molecular weight of 44 kDa was detected. It can be concluded that gD1 is a monomer in solution.

**Figure S 6: Sequence alignment of EHV-1 with HSV-1 and PrV.** Sequence alignment based on secondary structures of gD1 with (A) HSV-1 (PDB ID 2C3A) and (B) PrV (PDB ID 5X5V) gD according to dssp [72]. Sheets are indicated as pink arrows, helices as blue cylinder, disulfide bonds as yellow boxes, glycosylation sites in gD1 as green dots, and magnesium coordinating residues in gD1 as purple dots. Labels correspond to the naming scheme presented by Li et al. [29].

**Figure S 7: Ramachandran plots for modeled MHC-I.** Ramachandran plots for (A) MHC-I (gene 3.1) and (B) equine β2m. Symbol code: green point-residue with favorable geometry, yellow point- residue with allowed geometry.

